# Formation of an RNA-mediated nuclear compartment

**DOI:** 10.64898/2026.08.20.746005

**Authors:** Denis L. Lafontaine, Ezequiel Calvo-Roitberg, Yu-Chieh Chung, Liyan Yang, Nevin Wise, Li-Chun Tu, Athma A. Pai, Job Dekker

## Abstract

Mammalian nuclei are spatially compartmentalized so that active and inactive segments of the genome occupy separate sub-nuclear neighborhoods. Compartments are often found in association with nuclear structures such as the nuclear lamina, nucleoli, and nuclear speckles (speckles), suggesting links between them. The molecular mechanisms by which compartments form remain largely unknown. Speckles are nuclear bodies that contain high concentrations of RNA splicing factors and associated chromatin has a high density of highly expressed genes. Combining liquid chromatin Hi-C to quantify chromatin interaction lifetimes genome-wide, immunofluorescence, fluorescence *in situ* hybridization, live cell imaging, and nascent transcript analysis, we have determined the biophysical and molecular basis of the speckle-associated chromosomal compartment. We find that genomic regions making up this compartment are stably glued together. Surprisingly, removal of speckles by rapid depletion of SON and SRRM2 that form the structural scaffold of these bodies shows that this stable association is not dependent on the speckle itself. Instead, we find that RNA molecules are the molecular glue that forms the biophysical basis of the speckle chromatin compartment. Based on observations that promoters and enhancers are also engaged in stable long-lived chromatin interactions that are independent of RNA, we propose a pathway for formation of the speckle chromosomal compartment: Initial stable clustering of promoters and enhancers is followed by production of (nascent) RNA. The exceptionally high density of GC-rich RNA emerging from speckle-associated loci forms a glue that holds these loci together and facilitates recruitment of speckle components. The result is a structurally stable nuclear compartment that facilitates efficient splicing.

## Introduction

The mammalian cell nucleus is structurally and functionally compartmentalized. From a cytological point of view this is apparent at the level of the whole nucleus by the fact that transcriptional silent closed and condensed chromatin is mostly localized at the nuclear periphery, while active and open chromatin is located more centrally^1–6^.

Compartmentalization is readily detected by genomic assays such as Hi-C and SPRITE. These assays detect interaction frequencies between pairs (Hi-C^7^) or groups of loci (SPRITE^8^) and have been used to define compartments and sub-compartments based on their genome-wide interaction patterns. Globally, two types of major compartments can be defined that correlate with open and active chromatin (euchromatic A compartments), and closed and silent chromatin (heterochromatic B compartment). A compartment domains preferentially associate with other A compartment domains, and similarly, B compartments interact more frequently with other Bs. A and B compartments can be further split in sub-compartments based on their chromatin properties (e.g., histone modifications), their long-range chromatin interactions, and associations with nuclear structures such as the lamina^9–11^. For instance, the A compartment can be split in several types of active chromatin. As compartments, sub-compartments tend to interact most with other loci of the same sub-compartment status.

The nucleus also harbors a variety of membrane-less organelles, or nuclear bodies. These non-chromatin structures are often located at chromosome territory interfaces^12–16^. Examples of such nuclear bodies include nucleoli and nuclear speckles. These bodies are formed through condensate formation around some key scaffold components (e.g., the SON and SRRM2 proteins for speckles^17^) and contain high concentrations of reactants and substrates. Each is dedicated to specific nuclear processes: the nucleolus is the site of rRNA expression and processing (e.g., reviewed in^18^) and nuclear speckles are sites of splicing and mRNA processing^17,19^. Each of these nuclear bodies are surrounded by specific segments of the genome further contributing to spatial compartmentalization of the genome: nucleolus-associated domains associate with the periphery of nucleoli^20^, while speckles are surrounded by highly expressed speckle-associated domains^8,21–23^. Recent studies have demonstrated that nuclear speckles are not just passive reservoirs of splicing machinery, but that these structures are essential for appropriate splicing of the transcripts emanating from the loci that surround these speckles^22,24^, while lower concentrations of splicing factors are sufficient for proper processing of active genes not near these bodies.

There are strong correlations between sub-compartmentalization of chromosomes observed with Hi-C/SPRITE, and nuclear bodies and landmarks suggesting that their associations may play roles in structural and functional compartmentalization. For instance, several B type sub-compartments are anchored at the nuclear periphery through association with lamins and other proteins such as the lamin B receptor. Similarly, a specific A sub-compartment (“A1” as defined by Rao et al and Spracklin et al.^9,10^, and as a “Speckle” sub-compartment by the SPIN approach^11^. A1 or “Speckle” domains are enriched in highly expressed GC-rich genes clustered around nuclear speckles. Most genes typically contain long AT-rich intronic regions and more GC-rich exons. In contrast, genes associated with nuclear speckles are distinct: they possess introns and exons that have similar or “leveled” sequence composition^22,25,26^.

Interestingly, the appearance of domains enriched in “leveled” genes also evolutionarily coincides with the appearance of nuclear speckles as organisms transitioned from aquatic to terrestrial life^22^. It is possible that this unique intron-exon organization requires the high concentrations of splicing and RNA processing factors offered by close association with nuclear speckles to drive the very high expression and distinct splicing requirements of these genes.

Although compartmentalization of the nucleus is well known and increasingly understood to be functional (e.g., as for nuclear speckles and associated speckle sub-compartment), the mechanism driving (sub-)compartmentalization of the genome are much less understood. One model is that chromosome compartmentalization is driven by a process of phase separation. In this model sub-compartment domains display homotypic affinities for each other, and possibly with non-chromatin structures and nuclear bodies. Such a system will tend to phase separate with sub-compartment domains of different types segregated from each other. From a polymer theory point of view such a system has been shown to be well approximated as a block co-polymer, and such models can reproduce Hi-C data and images of compartmentalized nuclei^7,27–29^. Phase separation of segments of chromatin of different types, as defined by histone modification patterns, has also been directly observed in vitro^30^, adding further support for this mechanism. A key biophysical driver of phase separation is differential affinities between pairs of loci: phase separation will only occur when affinities between pairs of the same sub-compartment are higher than affinities between pairs belonging to different sub-compartments ^31–33^. However, the identities of the components and molecules of chromatin that determine these affinities are often not known, though roles for histone modifications^30^ and nuclear bodies^34^ have been suggested.

Previously we developed “liquid chromatin Hi-C”, or LC-Hi-C^29^. LC-Hi-C is a Hi-C variant that can measure the lifetime of chromatin interactions genome-wide. Briefly, in LC-Hi-C, chromosomes are restriction-digested *in situ*, followed by incubation for up to several hours to allow loci interacting through weak or dynamic interactions to dissociate, leaving only stable interactions. Cells are then fixed, and Hi-C is used to identify the stable interactions that remain and, by comparison to undigested chromosomes, the weak or transient interactions that were lost. We found that A-compartment chromatin is generally more dynamic than that in the B-compartment^29^.

Here, we aimed to identify molecules that can mediate stable long-lived interactions between sub-compartment domains with the intent to learn more about general mechanisms of nuclear and chromosomal compartment formation. We leveraged the ability to measure chromatin interaction lifetimes using LC-Hi-C to identify loci with stable interactions with other loci. We show that nuclear speckle-associated chromatin domains display among the most stable and long-lived chromatin interactions in the nucleus. We then go on to show through a series of perturbation experiments that this stability is dependent on RNA, and not on the association of chromatin with nuclear speckles, nor the structural integrity of the speckle itself. Further, kb-resolution analysis of LC-Hi-C data showed that pairs of promoters and enhancers in general are also engaged in long-lived interactions, but these are not dependent on RNA. Based on our observations we propose a pathway for the formation of the stable speckle-associated compartment.

## Results

### Speckle-associated chromatin interactions stabilize in a growth-dependent manner

Previously, we used LC-Hi-C on K562 cells and reported that chromatin interactions in active euchromatic regions are relatively short-lived whereas interactions in heterochromatic domains are longer-lived^29^. Here, we set out to explore the mechanistic basis of the relationships between chromatin conformational stability, chromatin state, and gene expression.

We performed LC-Hi-C on a culture of exponentially growing K562 cells. Nuclei were isolated and incubated for 2 hours in the presence of DpnII (DpnII-digested) or in restriction buffer without restriction enzyme (Mock-digested). Nuclei were then fixed and Hi-C was performed. We previously showed that chromosome conformations detected by Hi-C are strikingly similar in intact cells or in mock-digested isolated nuclei^29^. **Fig. 1a** shows a Hi-C interaction map for chromosome 11 obtained with mock-digested nuclei. This map displays the typical inverse correlation between interaction frequency and genomic distance, and a prominent checkerboard patterning reflecting compartmentalization, exactly as observed with intact cells. For DpnII-digested nuclei, however, the Hi-C maps change greatly (**Fig. 1b**): first, the overall interaction maps become less crisp; second, there is an increase in longer-range interactions (*in cis* and *in trans*^29^) leading to a less intense main diagonal; and third, a blurring of the checkerboard pattern. As we described before^29^, these changes in Hi-C interaction patterns represent liquification of chromatin, where interacting loci dissociate from each other upon digestion, become mobile, and move away from their initial spatial positions, resulting in loss of chromatin interactions.

**Fig. 1:**
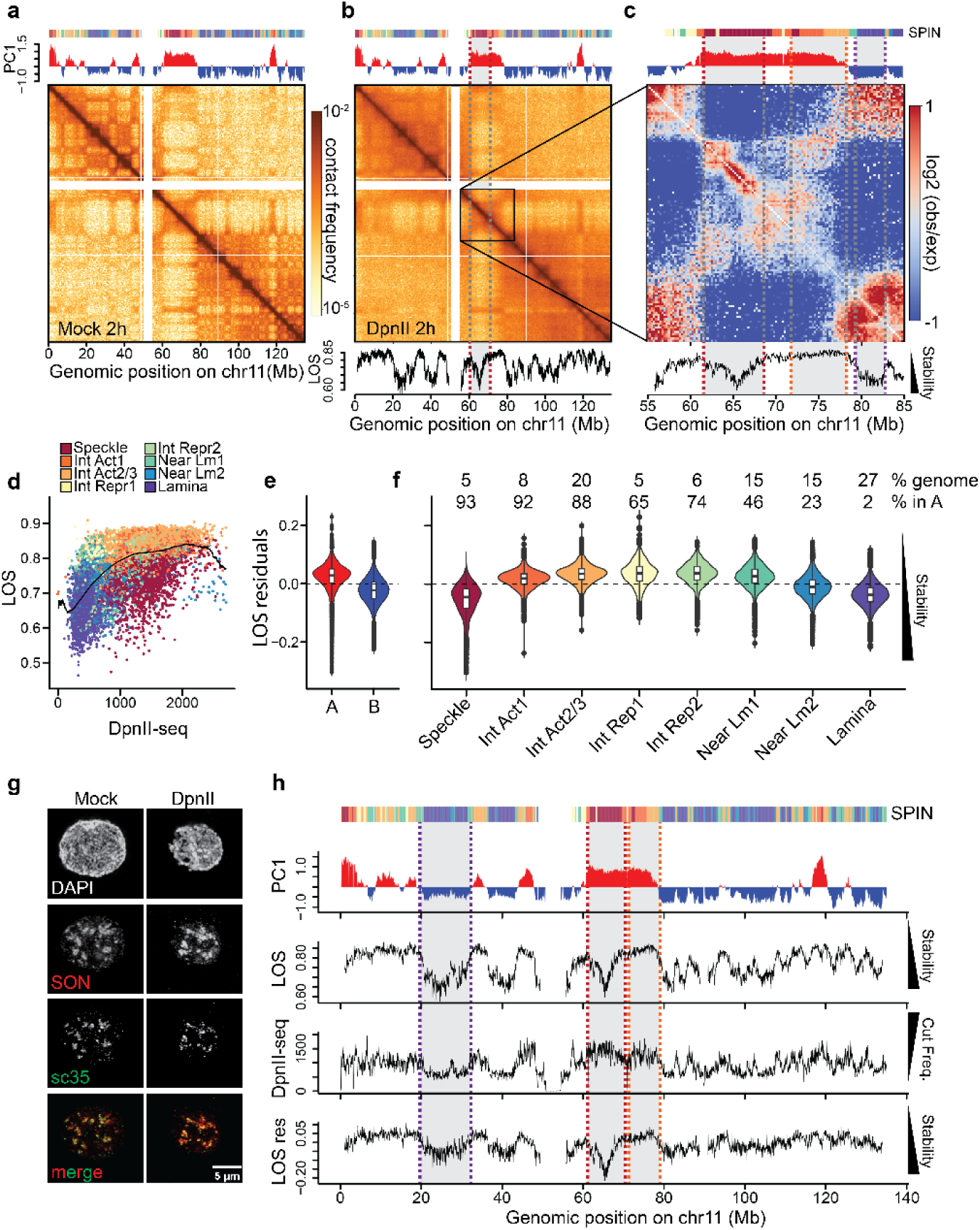
LC-Hi-C demonstrates that Speckle-and lamin-associated chromatin interactions display high stability. Heatmaps of chromosome 11 generated after isolated nuclei were (**a**) Mock-and (**b**) DpnII-digested for 2 hours before fixation and Hi-C. PC1 track generated by eigenvector decomposition of the mock-digested heatmap. SPIN annotations are depicted in the bar at the top (Speckle = Nuclear Speckle; Int Act1 = Interior Active 1; Int Act2/3 = merged Interior Active 2 and Interior Active 3; Int Repr1 = Interior Repressed 1; Int Repr2 = Interior Repressed 2; Near Lm1 = Near Lamina 1; Near Lm2 = Near Lamina 2; Lamina = Nuclear Lamina). LOS tracks are plotted below the heatmap. **c**, Zoom into an A compartment domain on chromosome 11 and plotting the log2(observed/expected) Hi-C contacts generated from the DpnII-digested Hi-C data. Lamin-(purple dotted line), speckle-(red dotted line) and non-speckle-associated active chromatin (orange dotted line) domains are highlighted. **d**, LOS data plotted against DpnII-seq signal (colored by SPIN state) to calculate the residuals from each LOS data point to a moving average (black line). **e**, LOS residuals binned at 50kb plotted by compartment. **f**, LOS residuals plotted by SPIN state. Here we also denote the percentage of bins annotated in the genome (% genome) as well as the percentage of bins that are located within the A-compartment (% in A) for each SPIN category. Boxplots show the median (center line), interquartile range (box), and 1.5×IQR whiskers, with outliers plotted individually. **g**, 3D projections of mock-and DpnII pre-digested nuclei stained with DAPI. SON and sc35. **h,** Tracks of chromosome 11 showing compartment status using PC1 with SPIN state annotations in the colored bar above along with LOS, DpnII-seq signal and resulting LOS residuals. Lamin-(purple dotted lines), speckle-(red dotted lines) and non-speckle (A-compartment)-associated (orange dotted lines) chromatin regions are highlighted. All data is binned at 50kb except for heatmaps which are binned at 250kb.

We quantify these phenomena using the previously described Loss Of Structure (LOS) metric^29^. LOS quantifies, for each genomic bin, the relative loss of shorter-range interactions (between pairs of loci separated by up to 2 Mb) and corresponding gain in longer-range and inter-chromosomal interactions after pre-digestion^29^ (Methods). LOS ranges between 0 and 1. Higher LOS values represent more dissolution of chromatin interactions (“unstable” short-lived interactions), while lower LOS values represent regions where interaction frequencies are less affected by pre-digestion with DpnII (“stable” long-lived interactions). As we showed before^29^, LOS varies along chromosomes (plot below the Hi-C interaction map in **Fig. 1b**) in correlation with A/B compartmentalization (as captured by the first eigenvector EV1^7^, plot on top of the Hi-C interaction map). B compartment interactions typically display higher stability (lower LOS) than those in the euchromatic A-compartment^29^ (higher LOS) (**Extended Data Fig. 1a**).

Unexpectedly, we noticed several instances where loci embedded within large A compartment domains displayed lower LOS than adjacent loci within the same compartment domain, i.e., they displayed more stable and long-lived interactions than expected for A-type loci. A prominent example on chromosome 11 is shown in **Fig. 1b**. When we correct the observed interaction heatmap for the expected distance decay in interaction frequency [Log2(observed / expected)], the differential loss of chromatin interactions at the diagonal is particularly accentuated and correlates with the LOS metric (**Fig. 1c**). This result implies that A-compartment domains can contain sub-compartments with distinct properties. To explore this further, we made use of subcompartment annotations (“SPIN states”^11^) that were previously generated for K562 cells. SPIN state annotations are obtained by integrating Hi-C data with data on subnuclear positioning of loci (e.g., near the nuclear lamina based on LaminB1-DamID/TSA-seq^23,35^, near nuclear speckles based on SON-TSA-seq^23,36^, and near the nucleolus based on 4XAP3-DamID^11^. Ten SPIN states are defined as follows: “Speckle”, Interior Active 1 (“Int Act1”), Interior Active 2 (“Int Act2”), Interior Active 3 (“Int Act3), Interior Repressed 1(“Int Repr1”), Interior Repressed 2 (“Int Repr2”), Near Lamin 1 (Near Lm1), Near Lamin 2 (“Near Lm2”), Lamina, and Lamina-like (**Extended Data Fig. 1b**). The Speckle and Int Act1 states are the most highly transcribed regions (**Extended Data Fig. 1c**), both displaying association with SON, a key structural component of speckles. However, they differ in the level of SON association, with Speckle loci having higher levels than Int Act1 loci, suggesting both are speckle-associated but with higher and lower frequency respectively. Int Act2 and Int Act3 SPIN states are also actively transcribed loci but display low SON association and have otherwise highly similar features. We therefore merged these two states into a group named “Int Act2/3”. The two Interior Repressed SPIN states are enriched in nucleolar markers but differ in SON and H3K27Me3 levels. Finally, the four Lamina SPIN states represent loci near the nuclear lamina, but with different enrichments of Lamin-DamID signal. The “Lamina-like” loci that can be uniquely mapped, e.g., in Hi-C, represent only ∼1% of the genome, often immediately adjacent to centromeres, and were left out of the analysis. When we plotted SPIN states along chromosome 11 (colored bars above the eigenvector tracks in **Fig. 1a, b, and c**, we noticed that the region with stable long-lived chromatin interactions embedded within the large A compartment domain corresponds to a sub-compartment domain annotated as “speckle-associated”.

LOS can be influenced by variation in DpnII pre-digestion bias along the genome. As in^29^, we employ DpnII-seq to first measure cut frequencies genome-wide and then use this information to correct the LOS metric for any such frequency bias. In this assay, nuclei are isolated and restriction enzyme digestion is performed. However, instead of fixation followed by Hi-C, the reaction is stopped, DpnII-digested ends are biotinylated and pulled down for sequencing (Methods). LOS correction is done by first plotting LOS as a function of DpnII-seq cut frequency for genomic bins, then calculating a moving average across the data and measuring the residual in LOS from each point to the average (**Fig. 1d**). Plotting LOS residuals by compartment status, we see that the overall differential stability between A-and B-compartments persists after digestion bias correction (**Fig. 1e**). If we instead subset by SPIN state^11^, we see that speckle-associated chromatin, which is largely located in the A-compartment (93%) and makes up only 5% of the K562 genome is very stable, i.e., displays low LOS (**Fig. 1f**; **Extended Data Fig. 1d**). Nuclear speckle morphology is largely preserved after DpnIII digestion (**Fig. 1g**). Note that, unlike lamin-associated heterochromatin which has lower cut frequency, speckle-associated chromatin is the SPIN state displaying the highest cut frequency as measured by DpnII-seq (**Extended Data Fig. 1e**). Yet, after correction for differences in DpnII-digestion, both display stable chromatin interactions (low residual LOS values). This phenomenon is seen along chromosome 11, where stable lamina-associated chromatin is cut to a lesser extent than A-compartment chromatin (**Fig. 1h**). However, A-compartment chromatin can vary widely in local stability, yet cut frequency remains relatively uniform throughout the A-compartment domain. Throughout this study, we performed matching DpnII-seq for each LC-Hi-C experiment, for all perturbations and replicates.

A subset of telomeres is known to associate with nuclear speckles while others interact with speckles in a cell cycle-dependent manner^37^. In our data we also see examples of speckle-associated domains near telomeres and these display high chromatin interaction stability (**Extended Data Fig. 1f**). Globally, we see that 34% of all speckle-associated regions fall within 10 Mb from telomeres and that 22/41 detectable (i.e., by sequencing and excluding telomeres from acrocentric arms) telomeres contain at least three 50 kb speckle-associated bins within 10 Mb of the chromosome end (**Extended Data Fig. 1g**). Other SPIN annotations near telomeres display similar stability as observed throughout the genome, indicating that proximity to a telomere does not stabilize chromatin interactions generally. On the other hand, other than the small and unique “Lamina-like” SPIN group, pericentromeric regions do not appear to be particularly enriched in any SPIN state (**Extended Data Fig. 1h**).

In our previous LC-Hi-C analysis in K562 cells we did not notice these very stable speckle-associated chromatin interactions^29^. Given that genomic regions associated with nuclear speckles are gene-rich and highly active, we hypothesized that growth conditions could contribute to the stability of interactions at these sites. To test this, we grew K562 cultures for 7 days without passaging or adding additional media. This treatment resulted in saturation of the culture without appreciable cell death, i.e., cells became quiescent but did not die (**Extended Data Fig. 1i**). LC-Hi-C data generated from these quiescent cultures show that 1) fragment size distributions between quiescent and control cultures were digested to the same extent (**Extended Data Fig. 1j**) and 2) speckle-associated chromatin displays a decrease in interaction stability, recapitulating the absence of speckle-associated stability observed in^29^ (**Extended Data Fig. 1k**). These observations demonstrate that growth conditions, especially quiescence, can dictate speckle-associated chromatin interaction stability and offer an explanation as to why stable chromatin interactions at speckle-associated regions were initially not observed^29^. Interestingly, during quiescence transcription levels are generally reduced^38^, suggesting a link between gene expression levels, speckle association, and chromatin interaction stability. Below we explore this directly.

### Perturbations in transcription and splicing destabilize speckle-associated chromatin interactions

Perturbations in RNA metabolism such as inhibition of transcription^39,40^ and splicing alter nuclear speckle morphology. Given these observations, we set out to test whether such perturbations lead to changes in chromatin interaction stability. We inhibited transcription by treating cells with 25 uM triptolide and 200 uM 5,6-Dichloro-1-β-D-ribofuranosylbenzimidazole (TPL/DRB) for four hours. In a parallel set of experiments, we blocked splicing by transfection of a U1 anti-sense morpholino (AMO) followed by a 4-hour incubation. Both control and U1 AMOs were tagged with fluorescein which allowed us to calculate the transfection efficiency, which was > 90% for all transfections. The use of fluorescein also allowed us to track the intracellular localization of the AMOs. We observed that the control AMO (cAMO) was uniformly diffused throughout the nucleoplasm whereas the U1 AMO colocalized with speckles as predicted (**Extended Data Fig. 2a**).

Both transcription inhibition and U1 inhibition treatments resulted in a change in the shape of speckles with speckles appearing rounder (**Fig. 2a**). Further, speckles increased in volume (**Fig. 2b**) and decreased in number, consistent with speckles fusing^40^ (**Fig. 2c**). It is known that nucleoli disorganize after inhibition of transcription^41^, which is confirmed in our data (**Fig. 2a**). However, nucleolar disorganization is not observed after U1 AMO treatment. Using SLAM-seq^42^, we observe a dramatic global reduction in total nascent transcripts (protein and non-protein coding genes transcribed by RNA polymerase II) after TPL/DRB treatment (10,029 of 12,765 genes are down regulated) and a less severe downregulation after U1 AMO treatment (6,395 of 12,481 genes are down regulated) (**Fig. 2d**). Very similar observations are made if we count reads that map only to exonic regions (**Extended Data Fig. 2b**). As expected, we see overrepresentation of intronic reads after U1 AMO treatment, but not after TPL/DRB treatment, indicating perturbations in RNA splicing (**Fig. 2e**). We conclude that U1 AMO-transfection inhibits splicing as well as transcription, consistent with earlier findings^43^.

**Fig. 2:**
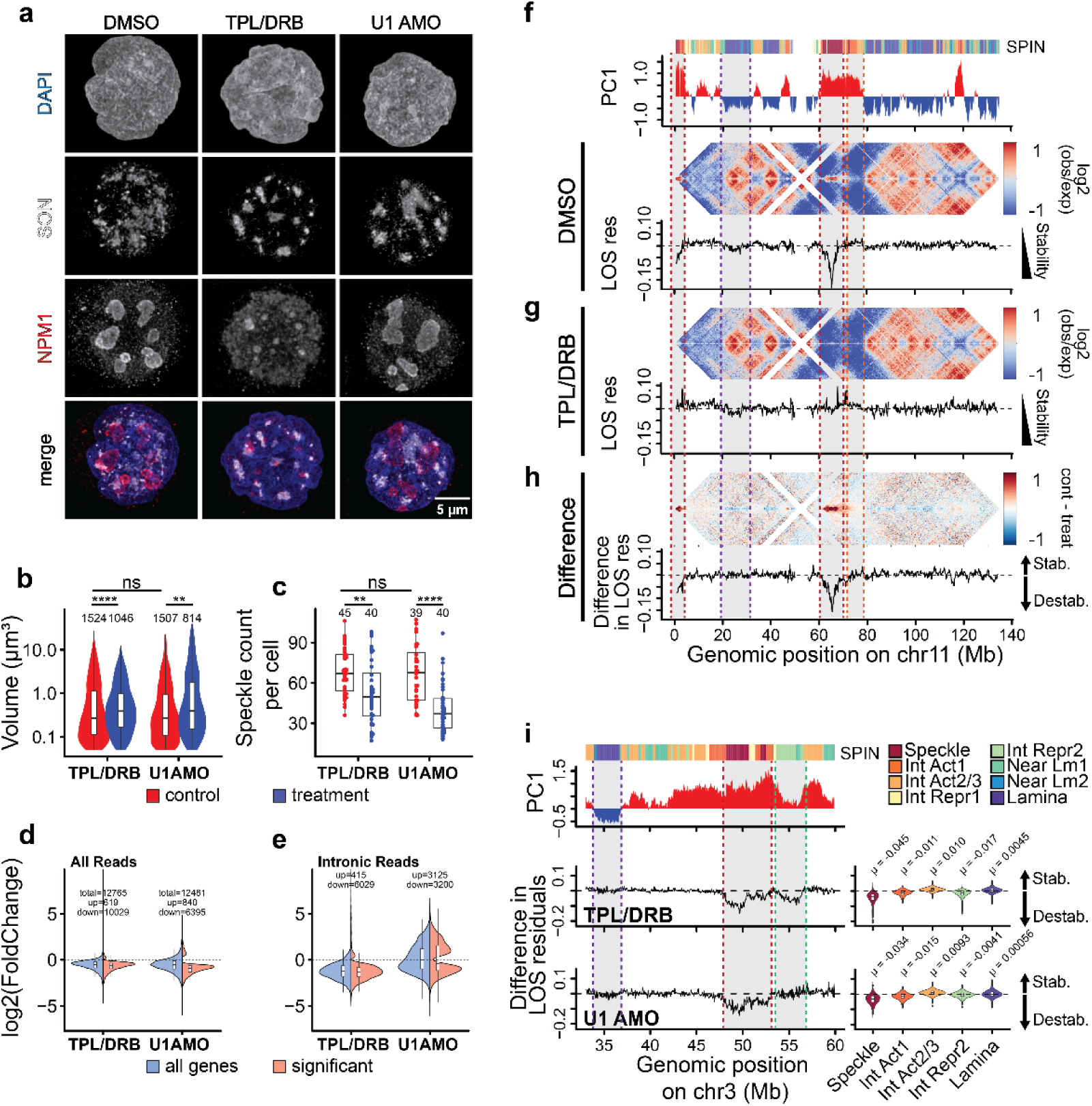
Disruption of nuclear bodies results in destabilization of associated chromatin interactions. **a**, 3D projections of cells stained with SON and nucleophosmin (NPM1) treated with either DMSO, triptolide/DRB (TPL/DRB) or U1 AMO for 4hours. Speckle volume (**b**) and speckle/cell (**c**) quantifications for at least 39 cells for both TPL/BRB and U1 AMO treatments with respective controls (DMSO = 45, TPL/DRB = 40, cAMO = 39, U1AMO = 40). Kolmogorov-Smirnov (K-S) tests were performed between control and treatment measurements (* p-value < 0.05, ** p-value < 0.01, *** p-value < 0.001). Significant Kolmogorov-Smirnov (KS) tests between DMSO vs TPL/DRB (p = 6e-28 and p = 0.003) and cont AMO vs U1 AMO (p = 0.005 and p = 0.000003) for both speckle volume and number, respectively, are observed. Differences in volume measurements and speckle counts between DMSO and control AMO were not significant (p = 0.32 and p = 0.08, respectively). Log2(FoldChange) values after differential gene expression analysis for ‘all genes’ (protein and non-protein coding) and ‘significant’ differentially expressed genes after TPL/DRB and U1 AMO treatments using all reads **(d)** and intronic reads **(e)**. **f**, log2(observed/expected) contact frequency heatmaps aligned with tracks of chromosome 11 showing SPIN states (bar above), PC1, and LOS residuals for DMSO treated cells. **g**, log2(observed/expected) contact frequency heatmaps and LOS residuals tracks for TPL/DRB treated cells. **h**, Heatmap and LOS residual track showing the difference between control (DMSO) and treated (TPL/DRB) log2(observed/expected) contact frequency heatmaps and LOS residuals, respectively. **i**, Tracks of chromosome 3 showing the difference in LOS residuals for TPL/DRB and U1 AMO treatments with their respective controls (control – treatment; left). Global difference in LOS residuals of bins within select SPIN state groups for each treatment (right). Global mean LOS values for each SPIN group are denoted. Bootstrap analysis generated significant p values (p < 0.0001) for comparisons of each SPIN group to 0. All boxplots show the median (center line), interquartile range (box), and 1.5×IQR whiskers, with outliers plotted individually. LOS tracks and SPIN violin plots are binned at 150kb. SPIN and PC1 tracks are binned at 50kb. Heatmaps are binned at 250 kb.

LC-Hi-C was performed after treatment with either TPL/DRB or the U1 and control AMOs. Pre-digestion of nuclei resulted in consistent fragment size distributions for each of the treatment/control pairs (**Extended Data Fig. 2c-d**). We first analyzed the effects of transcription inhibition on chromatin interaction stability. Visual inspection of the LC-Hi-C Log10(observed/expected) interaction maps for DMSO treated cells again shows the stability of speckle associated domains. For example, the region around position 60-70 Mb at chromosome 11 representing a major speckle-associated domain displays high observed/expected interaction values in LC-Hi-C maps, while flanking A-compartment loci display fewer interactions than expected indicating those interactions were unstable (**Fig. 2f**). Note that B-compartment domains also show elevated observed/expected interaction frequencies, as expected from the fact that these are also characterized by stable chromatin interactions. The LOS metric readily quantified the stability of the speckle-associated domain.

LC-Hi-C Log10(observed/expected) interaction maps obtained with nuclei isolated from cells after inhibition of transcription closely resemble those of control cells (**Fig. 2g**), with the notable exception that speckle-associated domains lost their frequent long-lived interactions. The LOS profile confirmed the loss of stable interactions at speckle-associated domains (**Fig. 2g**). To further quantitatively explore differences between TPL/DRB-treated cells and control cells we directly subtracted the corresponding LC-Hi-C Log10(observed/expected) interaction maps (**Fig. 2h**). In such a difference map, negative values correspond to destabilization and positive values correspond to stabilization of chromatin interactions after treatment. The difference LC-Hi-C map for chromosome 11 clearly shows several speckle-associated regions that display stable interactions in control cells and that are destabilized after transcription inhibition. This is also observed when we directly subtract the LOS residuals profile obtained with transcription-inhibited cells from the profile obtained with control cells (**Fig. 2h**). The strong dip in the difference in LOS residuals at position 60-70 Mb reflects the strong destabilization of chromatin interactions at that speckle-associated domain. We note the presence of another stable-speckle-associated domain at the left telomere that is also destabilized after transcription inhibition.

**Fig. 2i** shows another example of a stable speckle-associated region on chromosome 3. This region contains a speckle-associated domain embedded within a larger A compartment. The difference in LOS residuals (control – transcription inhibition) shows that chromatin interactions at the speckle-associated domain, but not at the flanking A compartment flanking at the left, are destabilized upon transcription inhibition. Interestingly, interactions at another domain to the right of the speckle-associated domain are also destabilized. This domain is annotated as “Interior Repressed 2”. This annotation includes nucleolus-associated chromatin and polycomb-associated loci (**Extended Data Fig. 1b**).

We then calculated the differences in LOS residuals (control – transcription inhibition treatment) genome-wide for each SPIN state separately (**Fig. 2i**, **Extended Data Fig. 2e**) and find that inhibition of transcription leads to the strongest destabilization of chromatin interactions at speckle-associated domains in general (Mean change in LOS residuals =-0.045) and to a lesser extent “Interior Active1” (Mean change in LOS residuals =-0.011) and “interior repressed 2” (Mean change in LOS residuals =-0.017) loci. “Lamina associated” chromatin interactions display the smallest changes in stability (Mean change in LOS = 0.0045).

Next, we analyzed the effects of U1 AMO-mediated transcription and splicing inhibition on chromatin interaction stability. LC-Hi-C data obtained with U1 AMO-transfected cells and analyzed in the same way as for transcription-inhibited cells, shows that chromatin interactions in speckle-associated domains are destabilized genome-wide (**Fig. 2i**, **Extended Data Fig. 2g**). The speckle-associated region on chromosome 3 shown in **Fig. 2i** is destabilized to the same extent when cells are treated with either the U1 AMO or transcription inhibitors. We note however, that some speckle-associated regions, notably the domain at chromosome 11 are destabilized to a lesser extent (**Extended Data Fig. 2f**). Genome-wide, the effect of U1 AMO-transfection was less pronounced than transcription inhibition (Fig. 2i; Mean change in LOS residuals: “Speckle”=-0.034, “Interior Act1”=-0.015, “Interior Repr2”=-0.0041, and “Lamina”= 0.00056). Interestingly, in contrast to what we observed with transcription inhibition, U1 AMO-transfection did not destabilize “Interior Repressed2” chromatin (**Fig. 2i** bottom, **Extended Data Fig. 2g**). The destabilization of Interior Repressed regions after transcription inhibition but not U1 AMO treatments, correlates with differences in nucleolar integrity observed by imaging after transcription inhibition and U1 AMO transfection (**Fig. 2a**).

Previous studies have reported that blocking transcription in K562 cells results in only very minor changes to chromatin conformation as observed by conventional Hi-C^44^. We examined and compared conventional Hi-C data generated with treated and untreated mock-digested nuclei. Here we also observe only subtle changes in the log2(obs/exp) contact frequency with conventional Hi-C along chromosome 11 (**Supplementary Fig. 1a**). For instance, the large speckle-associated domain at position 60-70 Mb displays a small <u>L</u>oss <u>O</u>f Hi-C <u>C</u>ontacts (LOC; calculated exactly as LOS but then applied to conventional Hi-C data) after transcription inhibition, but this effect is much smaller than the observed change in LC-Hi-C interactions.

Further, we see some attenuation in compartment strength globally, particularly in A and speckle-associated compartments (**Supplementary Fig. 1b,c**). Very similar observations were made when comparing control AMO and U1 AMO treated cells. We do see some local changes in log2(obs/exp) contact frequency along chromosome 11 (**Supplementary Fig. 1d**), however these changes are distinct from those observed with LC-Hi-C. Again, other than minor changes in compartment strength in A and speckle-associated chromatin, compartmentalization is generally unaffected (**Supplementary Fig. 1e,f**). When we performed the same analyses to quantify compartmentalization strength, e.g., using saddle plots based on PC1 or by direct pair-wise analysis between sub-compartments, using LC-Hi-C data we detected larger attenuation of compartmentalization strength: both in cis and in trans, interactions at and between speckle-associated domains were destabilized after transcription block, and to a lesser extent after U1 treatment (**Supplementary Fig. 1g-j**). This suggests that inhibition of transcription not only destabilizes interactions throughout speckle associated domains, but also between domains located on the same chromosome or on different chromosomes. LC-Hi-C detects changes in chromatin interaction stability that are not readily detected by conventional Hi-C that captures steady state chromatin conformation.

### Decrease in chromatin conformation stability correlates with decreased speckle association

Given that transcription and splicing inhibition cause both changes in speckle morphology and destabilizes chromatin interactions, we hypothesized that association of chromatin domains with nuclear speckles physically stabilizes their chromatin interactions, possibly via RNA. We should therefore be able to visualize direct contacts of speckle-associated chromatin domains with speckles, and these contacts should be lost after treatments such as transcription inhibition. To test this, we performed immunofluorescence (IF) microscopy using antibodies against SON coupled with DNA FISH labeling of specific loci. We selected an expressed locus that is destabilized after transcription inhibition (chr3_50M focus; **Extended Data Fig. 3a**) and a control locus that is also expressed but is not destabilized by transcription inhibition (chr15_75M; **Extended Data Fig. 3b**). Both these loci are annotated as “speckle-associated” based on SON-TSAseq and have similar gene density and expression levels. We find that the chr3_50M locus is frequently associated with speckles in control cells but is located farther from speckles after transcription inhibition (**Fig. 3a**). The chr15_75M locus also associates with speckles in control cells but to a lesser extent than the chr3_50M focus. To quantify these observations, we measured the amount of SON signal directly at and surrounding the two loci. Specifically, we first segmented z-sections containing labeled DNA foci and then created a dilated mask of 5 pixels surrounding the segmentation mask (**Extended Data Fig. 3c**). We measured SON intensity within both segmentation and dilated masks. First, we see that the chr3_50M locus generally has higher SON intensity (p-value = 2.8e-10; **Fig. 3b**) than the chr15_75M locus, despite producing smaller (**Extended Data Fig. 3d**) foci in images that display overall lower mean SON intensity (**Extended Data Fig. 3e**). This demonstrates that although both loci are annotated as speckle-associated, they differ in the extent to which they interact with speckles. Accordingly, these loci differ in their SON TSA-seq signal where the chr3_50M loci falls in a region with higher values than chr15_75M (**Extended Data Fig. 3f**). Second, the SON signal at and around the chr3_50M locus is significantly reduced after transcription inhibition, which is consistent with dissociation of these loci from speckles after treatment (p-value = 0.000026; **Fig. 3b**). No changes were observed in SON overlap with the chr15_75M locus before and after transcription inhibition (p-value = 0.057; **Fig. 3b**). These data show that transcription inhibition results in dissociation of speckles and their associated genomic loci, and dissociation from speckles correlates with destabilization of chromatin interactions. Loci with lower SON-association (i.e. the chr15_75M locus which displays less overall SON enrichment as measured by both SON TSA-seq and imaging) tend to have higher LOS in control cells and these loci do not further destabilize upon transcription inhibition.

**Fig. 3:**
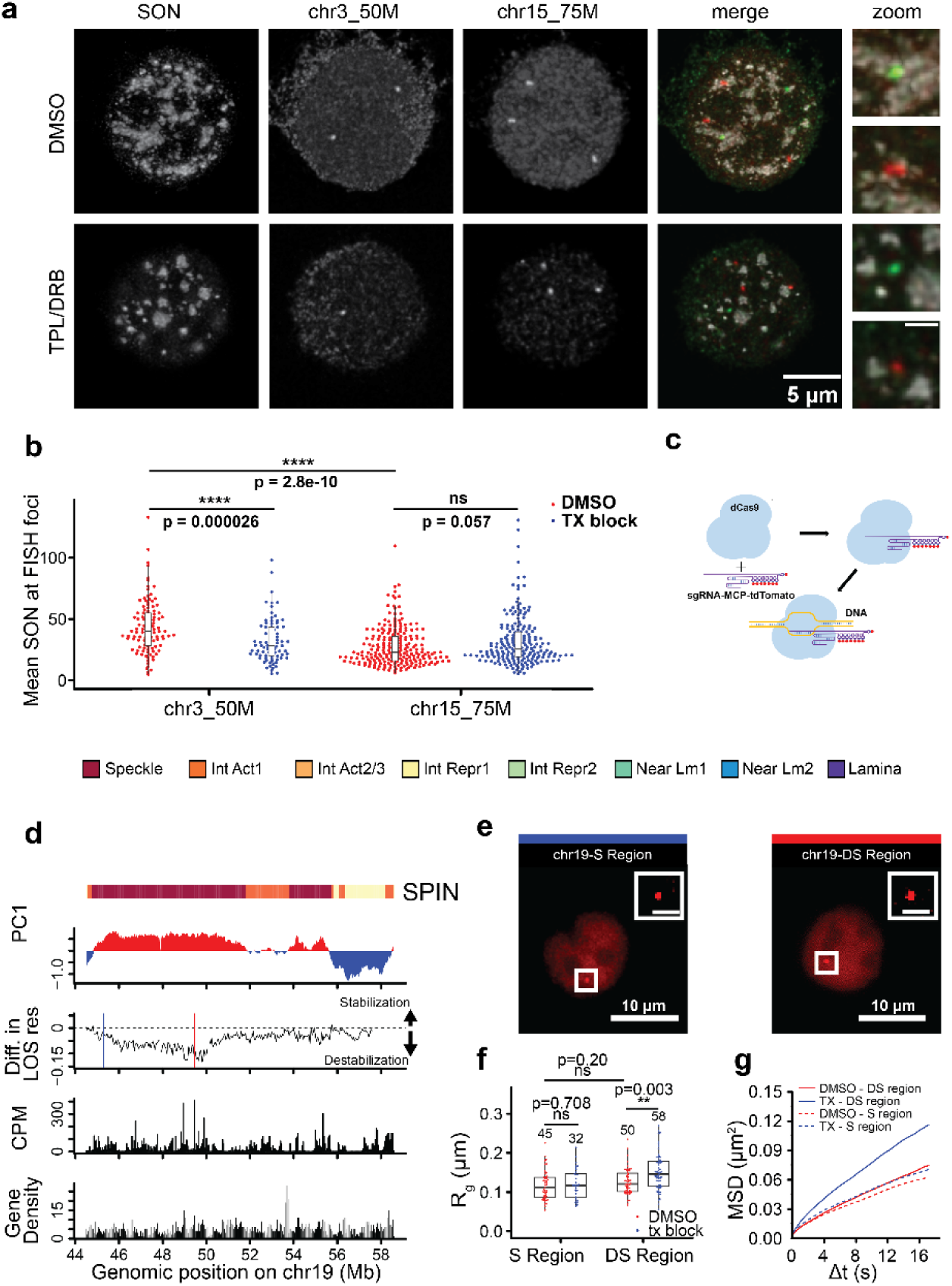
Destabilized loci show perturbed speckle association and increased mobility after transcription inhibition. **a**, Representative 3D projections where cells were treated with either DMSO or TPL/DRB, followed by fixation, staining with DAPI, SON antibody and probed with either the chr3_50M or the chr15_75M DNA-FISH probe sets. **b**, SON intensity was measured within and surrounding (5 pixel dilation) either chr3_50M or chr15_75M foci for at least 64 cells and 84 foci per treatment/probe combination (DMSO_chr3_50M = 109 foci / 71 cells, DMSO_chr15_75M = 226 foci / 89 cells, TPL/DRB_chr3_50M = 84 foci / 64 cell, TPL/DRB_chr15_75M = 202 foci / 80 cell). Kolmogorov-Smirnov (K-S) tests were performed between control and treatment measurements (* p-value < 0.05, ** p-value < 0.01, *** p-value < 0.001). **c**, A schematic of CRISPR-Sirius. The CRISPR-Sirius single-guide RNA (sgRNA) is engineered with an octet of MS2 hairpins (8×MS2). MS2 is labeled by binding to its coat protein (MCP), which is fused to the fluorescent protein tdTomato. **d**, Probes targeting SPIN annotated speckle-associated regions that either remain stable (blue line) or destabilize (red line) after transcription block were designed for live cell imaging. **e**, Tracks of target regions on chromosome 19 show SPIN state, compartment status, difference in LOS residuals after TPL/DRB treatment (control - treatment), CPM for nascent SLAM-seq reads and gene density. The genomic loci chosen for stable (chr19-S; blue line) and destabilized (C19-Destable, red line) regions of chromosome 19 in K562 cells. **f**, Box/swarm plots showing R_g_ values of loci in control stable and destabilized regions under DMSO and transcription-inhibition treatments. **g**, Ensemble-averaged MSD curves of loci in stable and destabilized regions under DMSO and transcription-inhibition treatments. Statistical significance was assessed using Welch’s t-test; *P < 0.05, **P<0.01, ***P < 0.005, and ns P > 0.05. All experiments were repeated more than three times. All boxplots show the median (center line), interquartile range (box), and 1.5×IQR whiskers, with outliers plotted individually. Genomic tracks are binned at 50kb.

### Decreases in chromatin conformation stability correlates with increases in chromatin mobility

Prior studies have shown that nucleosome dynamics in HeLa cells increase across the nucleus during transcription inhibition^45^, with similar increases observed for telomeres in HeLa cells and for loci in actively transcribed regions in U2OS cells^46,47^. We used CRISPR-Sirius imaging which previously has been utilized to visualize and track chromatin dynamics^47–50^, to assess if destabilization of chromatin interactions after transcription inhibition, as observed by LC-Hi-C, correlates with changes in chromatin mobility at these regions in live cells. In this assay dCas9 is loaded with gRNAs containing MS2 capsid protein binding sites to enable their visualization in K562 cells that express the MCP capsid proteins fused to tdTomato (K562-MCP-tdTomato in live cells (**Fig. 3c**). We designed two CRISPR-Sirius guide sets (chr19-S (<u>S</u>table) and chr19-DS (<u>D</u>e<u>S</u>tabilized); Supplementary Table 1, both located within a single large region annotated as speckle-associated on chromosome 19. The chr19-S region (**Fig. 3d**; blue vertical line) shows very minor changes in LOS whereas the chr19-DS region (**Fig. 3d**; red vertical line) is highly destabilized after transcription inhibition. We also selected these loci to be comparable in gene density and overall expression level in non-treated cells. We delivered sgRNAs and dCas9-P2A-HSA via electroporation (Methods, **Supplementary Fig. 2**). Both loci were readily localized, typically in the central part of the nucleus, as expected for highly active loci (**Fig. 3e**). After transfecting cells with the appropriate gRNAs, we treated cells with either DMSO or TPL/DRB for 4 hours and then imaged live cells at 0.2-second intervals (Δt=0.2 sec) for 85 consecutive frames. To characterize the dynamics of labeled loci within the stable and destabilized regions, we first obtained their trajectories by tracking real-time movements using 2D Gaussian fitting (**Extended Data Fig. 3g**), then used distances between point spread function centroids to calculate effective diffusion constants (D_eff_; **Extended Data Fig. 3h**), gyration radii (R_g_; **Fig. 3f**), and mean squared displacement (MSD) curves (**Fig. 3g**). D_eff_, obtained by fitting the MSD over a short time period (within 2Δt) using the normal diffusion equation, reflects short-time locus dynamics. The radius of gyration quantifies the spatial territory of locus movement, defined as the area explored by the locus (within 85Δt). MSD curves (within 85Δt) characterize long-time locus dynamics, and the dynamical parameters *D* and α can be extracted by fitting to the anomalous diffusion equation (Methods). Together, these parameters capture both short-and long-timescale behaviors of locus motion.

Without treatment (DMSO), the chr19-S and chr19-DS loci exhibited similar short-time dynamics having similar D_eff_ (**Extended Data Fig. 3h**) and radii of gyration (**Fig. 3f**) values. Over long timescales, the MSD curves remain similar, with values of (D_app_=0.00361, α=0.64267) and (D_app_=0.00326, α=0.70112) for chr19-S and chr19-DS, respectively (**Fig. 3g**; Supplementary Table 2). The D_eff_ values of loci in stable regions are unchanged (p = 0.16) after transcription-inhibition treatment. In contrast, D_eff_ values in the destabilized region significantly increase after transcription-inhibition treatment (p < 0.001; **Extended Data Fig. 3h**). R_g_ values of loci in stable regions remain similar after transcription-inhibition treatment (p = 0.708), whereas Rg values in destabilized regions increase significantly (p = 0.003; **Fig. 3f**).

Taken together, all metrics derived from CRISPR-Sirius live cell imaging support an increase in mobility of loci whose chromatin interactions destabilize after transcription inhibition as observed by LC-Hi-C. However, loci that display no change in interaction stability after transcription inhibition, do not show changes in chromatin mobility.

### Nuclear speckles do not physically stabilize chromatin interactions

Ilik et al. demonstrated that depletion of SON and SRRM2 was sufficient to disrupt speckles and cause speckle components to mislocalize^17^. Using an inducible degron system that can simultaneously and rapidly deplete SON and SRRM2 in K562 cells^51^, we tested whether the speckle nuclear bodies physically stabilize chromatin interactions. The addition of dTAGV-1 stimulates the degradation of proteins tagged with FKBP12^F36V^. Here we treated cells with either dTAGV-1 which stimulates the degradation of proteins tagged with FKBP12^F36V^ or the compound dTAGV-1-NEG which is a binding-competent but degradation-dead control for 6 hours (**Fig. 4a**). We confirmed efficient combined SON and SRRM2 depletion (“speckle depletion”) by imaging (**Fig. 4b**). Speckle depletion resulted in the downregulation of 126 genes when considering all reads (**Fig. 4c**) and 927 genes when considering only reads originating from exons (**Extended Data Fig. 4a**). Similar to splicing inhibition using U1 AMO (**Fig. 2e**) we observe overrepresentation of reads originating from introns after speckle depletion (**Fig. 4c**). However, the number of genes with splicing defects is much smaller than when splicing in general is inhibited with a U1 AMO (**Fig. 2e**). In addition, unlike splicing inhibition using the U1 AMO where both downregulation of genes and intron over-representation is distributed across the genome covering genomic regions of varying SPIN states, after speckle depletion gene down-regulation and splicing defects occur almost exclusively at speckle-associated regions, and to a lesser extent Int Act1 loci (**Fig. 4d,e**). This is evident when looking at several chromosomes containing speckle-associated domains (**Fig. 4f**). These data show that nuclear speckles provide sub-nuclear structures required for effective splicing of genes in speckle-associated domains only, as reported recently by others^22^, while RNA from most other genes can be spliced without close proximity to speckles.

**Fig. 4:**
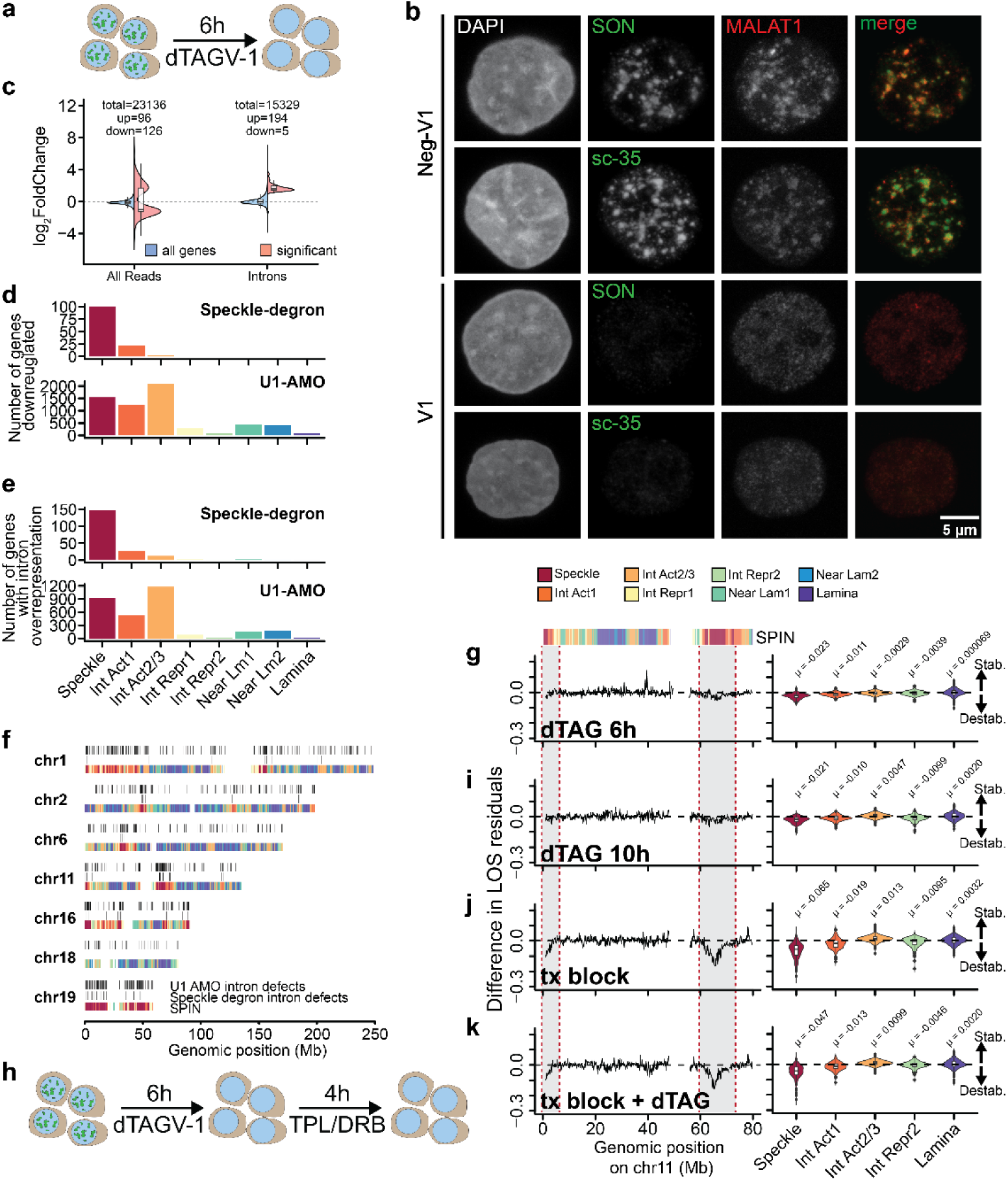
Speckle depletion/dissolution has little effect on chromatin interaction stability. **a**, Schematic of 6h SON and SRRM2 depletion strategy using the dTAG system. **b**, 3D Projections of fixed cells stained with either SON or SRRM2 antibody and probed with oligos against MALAT1 using RNA-FISH after treatment with dTag-V1 or dTAG-V1neg (control) for 6 hours. **c**, Log2(FoldChange) values after differential gene expression analysis for ‘all genes’ (protein and non-protein coding) and ‘significant’ differentially expressed genes after 6h SON and SRRM2 depletion for all reads and intronic reads. **d**, Number of down regulated genes by SPIN state for either SON/SRRM2 depletion or U1-AMO treatment. **e**, Number of genes displaying intron overrepresentation by SPIN state for either SON/SRRM2 depletion or U1-AMO treatment. **f**, SPIN state annotations across select full-length chromosomes showing sites that display intron overrepresentation after either SON/SRRM2 depletion or U1-AMO treatment. **g**, Tracks with SPIN annotations of chromosome 11 showing the difference in LOS residuals for 6h SON/SRRM2 depletion (left) and global difference in LOS residuals of bins within select SPIN state groups (right). **h**, Schematic of 10h SON and SRRM2 depletion using the dTAG system or 6h depletion followed by 4hrs TPL/DRB treatment. Tracks of chromosome 11 (left) and global differences in LOS residuals of of bins within select SPIN state groups (right) for 10h SON/SRRM2 depletion (**i**), 4h TPL/DRB treatment (**j**) and 6h SON/SRRM2 depletion followed by 4h TPL/DRB treatment (**k**). Bootstrap analysis generated significant p values (p < 0.0001) for comparisons of each SPIN group to 0. All boxplots show the median (center line), interquartile range (box), and 1.5×IQR whiskers, with outliers plotted individually. LOS tracks and SPIN violin plots are binned at 150kb.

We performed LC-Hi-C where cells were treated with either dTAG-V1 or dTAGv1-neg (control) for 6 hours (**Fig. 4a**). Pre-digestion of nuclei resulted in very similar fragment size distributions for each of the treatment/control pairs (**Extended Data Fig. 4b**). LC-Hi-C data demonstrate that speckle-associated regions again show high stability in this modified cell line (**Extended Data Fig. 4c,d**). Interestingly, we observe only a very modest destabilization of speckle-associated chromatin interactions after speckle depletion (**Fig. 4g**). While this minor genome-wide destabilization of speckle-associated domains is statistically significant, the change in stability is much smaller than what is observed in response to general transcription inhibition (**Fig. 4j**). The small change in interaction stability observed after speckle depletion may be related to the fact that speckle depletion results in down regulation of some genes at speckle associated domains (**Fig. 4d**). Further, given that splicing at speckle associated regions is defective after speckle depletion, while chromatin interaction stability is not changed to a large extent, it appears that splicing itself is not critical for stabilization of chromatin interactions.

We then tested whether speckle-associated loci in this cell line can be further destabilized by inhibiting transcription. Here, we subjected cells to either dTAG-V1 or dTAGv1-neg for 6 hours followed by either a DMSO treatment or TPL/DRB treatment for an additional 4 hours (**Fig. 4h**). First, we find that extending the speckle depletion from 6 hours to 10 hours does not further destabilize speckle-associated chromatin (**Fig. 4g,i**). Second, we find that transcription inhibition further and drastically destabilizes speckle-associated chromatin interactions in the presence or absence of nuclear speckles (**Fig. 4j,k**). These results indicate that transcription/RNA metabolism and not the speckle nuclear bodies themselves stabilizes chromatin interactions at and between speckle-associated loci.

### Further evidence that integrity of speckles is not required for chromatin interaction stability

Several observations suggest that nuclear bodies physically stabilize associated chromatin interactions: Perturbations in nuclear speckle morphology correlate with destabilization of speckle-associated interactions. Also, loci annotated as “Interior_Repressed”, covering nucleolus-associated loci, are destabilized after TPL/DRB but not U1 AMO treatments (**Fig. 2i**). Consistent with a physical nuclear body model, transcription inhibition, but not splicing inhibition, leads to disorganization of nucleoli (**Fig. 2a**). However, several lines of evidence argue against this physical interaction model. First, we tested whether heat-shock, which is known to perturb speckle morphology (**Extended Data Fig. 5a,b**), destabilizes speckle-associated chromatin conformation similar to that observed after TPL/DRB and U1 AMO treatments. Others have reported extensive down-regulation of transcription genome-wide after heat-shock using PRO-seq in K562 cells^52^. We exposed K562 cells for 80 minutes to 42 centigrade followed by imaging of speckles, LC-Hi-C, and nascent transcript analysis. In our experiment we observe fewer significantly differentially expressed genes overall (**Extended Data Fig. 5c**). Importantly, GO term enrichment for the significantly up-regulated genes (53) identified sets of genes related to heat-shock and protein refolding (**Extended Data Fig. 5d**), indicating that cells responded to the elevated temperature by expressing expected sets of genes. Speckle morphology changed as they rounded up (**Extended Data Fig. 5b**), similar to what is observed after transcription inhibition, and as reported previously^36,53^ (**Fig. 2a**). Fragment size distributions generated from pre-digestion of nuclei for LC-Hi-C were very similar between control and treatment for each perturbation (**Extended Data Fig. 5e**). Importantly, no striking differences in LOS residuals are observed between control and heat-shocked cells (**Extended Data Fig. 5f**).

Second, we showed that growth conditions can alter stability at speckle-associated regions (**Extended Data Fig. j,k**), in the absence of visible perturbations in speckle morphology (**Extended Data Fig. 5g**). Taken together, these results are not consistent with the model that nuclear bodies physically stabilize chromatin interactions along and between associated chromatin domains.

### Chromatin interaction stability at speckle-associated loci scales with transcription defects

Our results so far show that RNA metabolism, rather than the integrity of the nuclear speckle, stabilizes chromatin interactions at speckle-associated regions. In fact, we see that the difference in LOS residuals at speckle-associated regions negatively correlates with the number of genes that are down-regulated both genome-wide (**Fig. 5a**) and specifically within these regions (**Fig. 5b**). Thus, the larger the number of down regulated genes, the more the regions become destabilized. However, even though we observed a substantial number of downregulated genes within weakly speckle-associated (Interior_Act1) and non-speckle-associated active regions (Interior_Act2/3 SPIN states) after transcription block and U1-AMO treatment, no or only minor (for Interior_Act1) destabilization at these regions is observed (**Fig. 5c**, **Fig. 2i**). Further, destabilization in chromatin interaction stability due to transcription inhibition is restricted to the most highly expressed regions of the genome (**Extended Data Fig. 6a**).

**Fig. 5:**
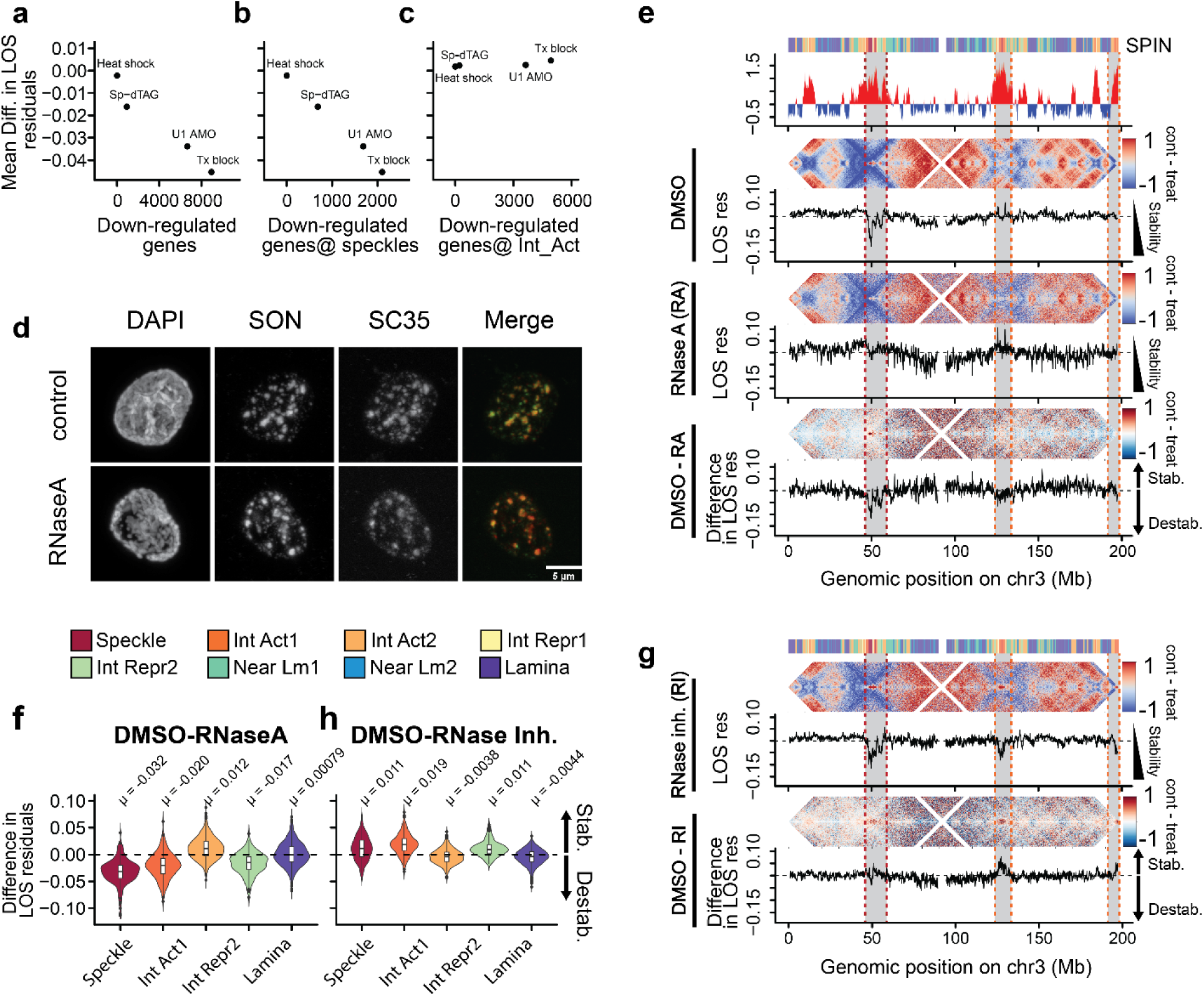
RNA stabilizes chromatin interactions at speckles. Scatter plots showing the mean difference in LOS residuals (binned at 50kb) at speckle-associated loci against the number of down regulated genes after TPL/DRB, U1-AMO, heat-shock treatments and SON/SRRM2-depletion genome-wide (**a**) or exclusively at speckles (**b**). **c**, Difference in LOS residuals (binned at 50kb) at Interior Act2/3 loci against the number of down regulated genes after TPL/DRB, U1-AMO, heat-shock treatments and SON/SRRM2-depletion at these regions. **d**, 3D projections of mock-or DpnII-digested nuclei treated with or without RNase A before staining with SON and sc-35 antibodies. **e**, Tracks for chromosome 3 showing PC1, SPIN annotations along with log2(observed/expected) heatmaps and associated LOS residual tracks generated from DpnII-digested control (top) and RNaseA-treated nuclei (middle). The difference between control and RNAse A-treated heatmaps and LOS residuals (bottom). **f**, Global difference between control and RNaseA-treated LOS residuals for select SPIN states. **g**, Log2(observed/expected) heatmap and associated LOS residual track generated from DpnII-digested nuclei isolated and pre-digested in the presence of RNase inhibitor (top) along with the difference between control and RNAse inhibitor-treated heatmaps and LOS residuals (bottom). Bootstrap analysis generated significant p values (p < 0.0001) for comparisons of each SPIN group to 0. All boxplots show the median (center line), interquartile range (box), and 1.5×IQR whiskers, with outliers plotted individually. LOS tracks and SPIN violin plots are binned at 150kb. SPIN and PC1 tracks are binned at 50kb. Heatmaps are binned at 250 kb.

Interestingly, high expression alone does not seem to be the sole determinant of the observed high stability as we observed a range of sensitivity to transcription inhibition at regions harboring high expression. The observed sensitivity of speckle-associated chromatin interaction stability to transcription inhibition at highly expressed regions correlates with proximity to nuclear speckles (as reflected in SON-TSAseq data) (**Extended Data Fig. 6b,c**) which, in turn, correlates with GC content (**Extended Data Fig. 6d**). Taken together, these data suggest that chromatin interaction stability at transcriptionally active regions is correlated with high transcriptional output, proximity to nuclear speckles, and GC content.

### RNA stabilizes chromatin conformation at speckle-proximal domains

The correlation of transcriptional output with chromatin interaction stability at speckles suggests that RNA itself stabilizes chromatin interactions. We therefore set out to test whether RNA molecules stabilize chromatin interactions. To do this we performed LC-Hi-C after isolated nuclei were depleted of RNA using 100 ug/ml RNase A (**Extended Data Fig. 6e**). DpnII pre-digestion of RNA-depleted nuclei resulted in very consistent fragment size distributions for each of the treatment/control pairs (**Extended Data Fig. 6f**). Although some morphological changes to chromatin are apparent when visualizing RNAse A treated DAPI-stained nuclei, as described before^54^, little changes are observed in the localization of speckle antigens (SON, SC35) after RNase A treatment (**Fig. 5d**). LOS analysis of LC-Hi-C data showed that RNAse A treatment destabilizes chromatin interactions at speckle-associated domains, consistent with a key role for RNA in gluing these loci together (**Fig. 5e,f**; **Extended Data Fig. 6g**, **Supplementary Fig. 4**). Interior Repr. 1 and 2 states, which represent loci near the nucleolus, also became destabilized (**Fig. 5f**; **Extended Data Fig. 6g**), consistent with what was observed when transcription is blocked (**Fig. 2i**; **Extended Data Figure 2e**).

The observation that RNA is key to interaction stability, made us wonder whether endogenous RNases modulate chromatin interaction stability in isolated control nuclei. To assess this, we included RNase inhibitor to nuclei extraction and pre-digestion steps followed by LC-Hi-C. Interestingly, the addition of RNase inhibitor increases the stability predominantly of Interior_Act1-associated regions and to a lesser extent, speckle-associated chromatin compared to control samples where RNase inhibitors were not included (**Fig. 5g,h**; **Extended Data Data Fig. 6h,i**). This suggests that RNA within Interior_Act1-associated regions also stabilize chromatin interactions, and that this was underestimated in nuclei isolated without RNAse inhibitors due to activity of endogenous RNases. As indicated above, Interior_Act1-associated regions display the second highest level of SON association, after loci annotated as “speckle-associated” (**Extended Data Fig. 1b**). Together, these results show that RNA is critical for chromatin interaction stability at speckle-associated regions, and at a set of additional regions near speckles (included in the SPIN category Interior_Act1-associated regions), but not at other active regions that are not at or near speckles (Int Act2/3-associated regions; **Extended Data Figure g,h**), or at inactive loci including lamina associated chromatin.

### Active cis regulatory elements show RNA-independent high chromatin interaction stability

Finally, we explored how chromatin interaction stability related to the presence of cis regulatory elements. We used SCREEN^55^ to retrieve genome-wide annotations of candidate cis-regulatory elements (cCREs) in K562 cells generated by the ENCODE project. We then calculated aggregated LOS at and around these elements. By combining multiple LC-Hi-C datasets from control conditions we were able to obtain sufficient read coverage to calculate LOS at 5 kb resolution. We then generated separate stack-up plots of LOS values (mean LOS residuals, as described above) centered at each type of cCRE (**Fig. 6a**, Methods). We find that LOS curves at active promoters and promoter-proximal enhancers (pELS, proximal enhancer-like sequence) display prominent dips, while other cCREs, including CTCF-bound sites, did not show such pattern. Proximal enhancers are defined as enhancers located within 2 kb of promoters, and thus in our analysis tend to be located within the same 5 kb bin as promoters. Examples of local LOS dips at individual active promoters are shown in **Extended Data Fig. 7a-c**. This suggests that active promoters and/or their nearby enhancers are engaged in relatively long-lived chromatin interactions with flanking chromatin.

**Fig. 6:**
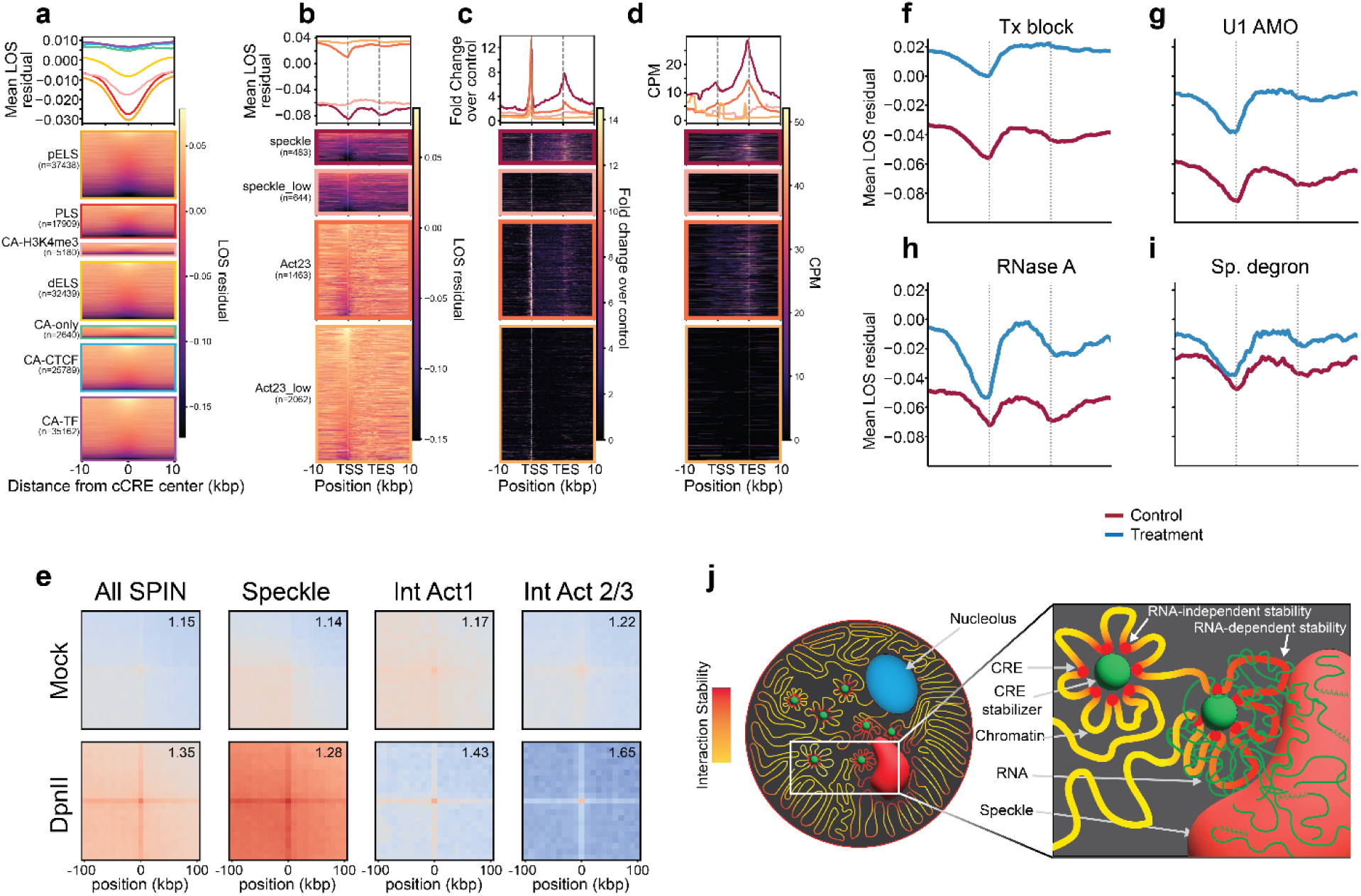
RNA-independent cCRE interaction stability. **a**, LOS residuals (binned at 5kb) centered on cCREs regions are shown individually and averaged by 200bp bins across the region. **b**, Scaled gene body regions of dominant transcripts showing LOS residuals anchored on TSS and TES are shown individually and averaged by 200bp bins. Gene bodies are divided by whether they belong to speckle-associated regions, Interior Act2/3 region each further subdivided by expression. Scaled gene body regions by SPIN state and expression showing PolII ChIP-seq (**c**) and spike-in normalized nascent RNA (**d**) signal where rows are sorted as in (b). **e**, 2D average of observed over expected interaction heatmaps for PLS pairs within 1 Mb for mock-and DpnII-digested Hi-C libraries, subset by Active SPIN states. Average LOS residuals across scaled gene bodies for control and treatment samples from the TPL/DRB (**f**), U1-AMO (**g**), RNaseA (**h**) treatments and SON/SRRM2 depletion (**i**) experiments. **j**, Model for RNA-dependent and independent mechanisms of chromatin interaction stabilization.

Next, we examined how chromatin interaction stability at active promoters and proximal enhancers depends on whether the gene is located within a speckle-associated domain or not. We plotted averaged LOS along 4 sets of scaled genes: active (TPM >5) and inactive genes (TPM = 0) embedded within speckle-associated domains or embedded within Interior Act2/3 domains that are not speckle-associated (**Fig. 6b**). We observe that the overall level of LOS along the gene and up-and downstream of the gene depends on whether genes are speckle-associated or not, as expected: LOS is lower for speckle-associated loci, i.e., active speckle-associated genes are engaged in longer-lived interactions. At a finer scale LOS dips at promoters and 3’ ends of genes of active but not of inactive genes, and this is observed for speckle-associated genes and for genes that are not speckle-associated. This is confirmed by plotting aggregated LOS plots at and around cCREs located in speckle domains or interior active2/3 domains (**Extended Data Figure 7d,e**): active promoters and proximal enhancers diaply local dips in LOS, while other cCREs do not, regardless of whether elements are located in speckle associated domains. We conclude that relative stability of chromatin interactions at promoters, and 3’ ends of genes, is a general phenomenon for active genes regardless of whether genes are speckle-associated.

We were interested to see what chromatin features along genes correlate with the LOS profile. We calculated average profiles along the same 4 categories of genes for RNA polII binding and nascent transcript levels (**Fig. 6c,d**). RNA polII levels correlate strongly with LOS, while nascent RNA levels do not suggesting RNA is not involved in stabilizing interaction at promoters.

To directly observe long-range interactions between pairs of active promoters (and adjacent pELS contained within the same 5 kb bin) we calculated averaged Hi-C and LC-Hi-C data for active promoter-promoter interactions (separated by up to 1 Mb) genome-wide, for pairs located within speckle-associated domains, Interior Active1, and Interior Active2/3 SPIN states (**Fig. 6e**). We observed enriched Hi-C and LC-Hi-C interactions between pairs of promoters. Importantly this is observed for all SPIN states, confirming these interactions are a general feature of active promoters, and not limited to speckles. In addition, while elevated promoter-promoter interactions are observed in Hi-C data, they stand out much more in LC-Hi-C data indicating that these interactions are longer lived than interactions between pairs of non-promoter interactions flanking these elements. Examples of stable local pair-wise interactions between individual active promoters are shown in **Extended Data Fig. 7a-c**.

Together these observations suggest that stability of chromatin interactions around active genes is determined at the domain level by proximity to speckles which is mediated by RNA, and at the cCRE level through additional other mechanisms. To test this, we plotted LOS around scaled active genes in speckle-associated domains before and after treatments that altered speckles and chromatin interaction stability at speckle-associated loci. After transcription inhibition with TPL/DRB (**Fig. 6f**) or through transfection of U1 AMOs (**Fig. 6g**), the overall LOS level throughout and flanking the genes was increased, i.e., chromatin interactions throughout the domains were destabilized as expected. However, the local dip in LOS at promoters and 3’ ends was retained. Similar results were observed after nuclei were treated with RNAseA (**Fig. 6h**), and to a much lesser extent in nuclei isolated after speckle depletion (**Fig. 6i**). We conclude that chromatin interaction stability at active cCREs is RNA, transcription, and speckle-independent, and thus represents a separate phenomenon from chromatin interaction stabilization at speckle-associated domains through RNA.

## Discussion

We identify RNA as a mediator of long-lived chromatin interactions at speckle-associated chromatin domains. We show that removing RNA through inhibition of RNA polymerase II or by RNAse A treatment destabilizes chromatin interactions throughout and between these domains. However, removing speckle scaffold proteins, which disrupts speckles and results in their components, e.g., the structural RNA MALAT1 (**Fig. 4b**) and speckle proteins SRRM1 and SF3A2^22^ to diffusely localize throughout the nucleus, did not change stability of chromatin interactions at and between these domains. This surprising result shows that stable chromatin interactions at and between speckle-associated domains are not the result of speckle association. Thus, while formation of the non-chromatin nuclear speckle body at these loci is essential for splicing of transcripts encoded by these speckle-associated domains, this structure is not involved in stabilizing interactions between these loci. We propose that RNA mediates the formation of a structurally stable chromatin compartment at which the non-chromatin speckle nuclear body is then assembled.

We find that RNA stabilizes chromatin interactions throughout speckle-associated domains, which can be hundreds of kilobases in size, including non-transcribed regions (**Fig. 6**). Therefore, we propose that RNA forms a general “glue” that spreads out throughout speckle-associated domains, extending beyond gene bodies, and that holds these domains together even after the chromatin is digested in small fragments of several kb during the LC-Hi-C procedure. Possibly the source of this RNA are the resident genes that are highly expressed leading to a high local concentration of RNA. Consistent with this idea is the observation that poly(A)+ RNA-FISH shows strong colocalization of the brightest poly(A)-RNA foci at speckles, indicating that these regions contain the highest local concentrations of poly(A)-RNA in the nucleus^22^. Other sources of RNA are also possible, e.g., RNAs transcribed elsewhere but that become localized at these domains. For instance, speckles contain non-coding RNAs such as MALAT1^19,21^, and these may associate with speckle-associated domains throughout the genome^56–58^. Finally, other types of RNA can be found associated with chromatin, including enhancer RNAs, upstream antisense RNA, a variety of repeat-generated RNAs (e.g., LINEs and SINEs), and non-coding RNAs involved in splicing, mRNA processing and modification. Currently we do not know the identity of the RNAs that mediate chromatin interaction stability, except that they are most likely generated by RNA polymerase II (given that inhibiting RNA polymerase II destabilizes chromatin interactions). Further, it seems less likely that the non-coding RNA MALAT1 is critical for this phenomenon: in cells in which we depleted speckles, MALAT1 became diffusely located throughout the nucleus and no longer enriched at specific regions (e.g., speckle domains), yet chromatin interaction stability at speckle domains was not reduced.

How could RNA act as a glue holding chromatin together? Nascent RNAs likely will recruit splicing factors and other RNA processing factors. It is possible that such factors can interact with multiple RNAs, effectively cross-linking them into a network that surrounds these highly expressed genes, possible via formation of condensates entrapping the chromatin. RNA driven condensates have been identified in other cases, including in the formation of the nucleolus^59,60^. It is also possible that RNAs directly hybridize with other RNAs, or that bridging RNAs are present. When a sufficiently high level of inter-linked RNA is present and these RNAs are also bound to chromatin, nearby chromatin segments become effectively glued together. We find that RNA specifically stabilized chromatin interactions at speckle associated domains, and to a lesser extent at domains annotated as “Interior Active 1” which contain loci that are also near speckles. These tend to be the highest expressed genomic loci. It is possible that what allows RNA to become a glue is simply the expression level: a certain amount of RNA may be required for this phenomenon to occur, but not all highly active genomic regions display chromatin interaction stability (**Extended Data Figure 6a**). It is possible that the nature of the RNAs is important. In that context it is intriguing that RNAs encoded by these regions are distinct from other active genes in the genome: these transcripts are more GC-rich and exon and intron lengths are “leveled”. It is possible that these RNAs have unique biophysical properties or binding partners. Finally, we note that our data are consistent with models proposed by Quinodoz and co-workers who also reported that classes of specific types of RNA correlate with clustering of chromatin into different types of compartments including speckles, the nucleolus, and pericentromeric heterochromatin^61^. Our data shows that in terms of mediating long-lived chromatin interactions, RNA appears to play a dominant role for the speckle compartment only, with much smaller effects for only one other compartment: chromatin associated with the nucleolus (**Fig. 5f**).

How do speckle-associated domains become associated with the non-chromatin nuclear speckle body? It is still an open question whether active genes move to speckles, or whether these regions recruit speckle components that then form a larger nuclear body through phase separation, through a “seed, bind and recruit” model^61^.

Recently a role for Alu RNAs has been proposed in mediating gene-speckle interactions, and it was proposed that these RNAs can form R-loops at target genes and recruit speckle components^57^. Several proteins, including p53^62^, CTCF, RAD21^63^, and MAZ^64^ have been implicated (reviewed in^19^). Belmont and co-workers have identified DNA elements that bind specific transcription factors that appear to mediate locus-speckle interactions, in some cases independent of transcription^65^. The latter observation provides further support for our data that the mechanisms by which chromatin interactions for speckle-associated domains are stabilized, which depend on transcription, can be separated from the mechanisms by which these loci associate with the speckle itself.

We also describe other types of chromatin interactions that are relatively long-lived but do not depend on RNA for interaction stability. First, as we described before^29^, lamin-associated heterochromatic domains are characterized by long-lived chromatin interactions, and this stability was not altered in any of the perturbations we performed here. Second, we find that promoters and proximal enhancers of active genes, regardless of whether they are located within speckle associated domains or not, are also engaged in RNA-independent and transcription-independent chromatin interactions with flanking chromatin and each other. We recently described such transcription-independent interactions between promoters during telophase using conventional Hi-C^66^. We found that these interactions transiently form during mitotic exit and then become either pruned by cohesin-mediated loop extrusion or these initially small promoter regions become resorbed within larger sub-compartments, especially the speckle sub compartment, and become more difficult to detect with conventional Hi-C^66^ (**Fig. 6e**). Importantly, we showed that these interactions are independent of transcription as they occur during telophase before transcription re-starts. Furthermore, these interactions became increasingly strong in cells with defects in nuclear transport which led to the exclusion of RNA polymerase II from the nucleus, further confirming they are independent of transcription. LC-Hi-C now shows these interactions stand out in normal interphase cells as compared to other nearby chromatin interactions by being more stable. What stabilizes these interactions is currently not known.

All together this led us to speculate how the nuclear speckle compartment is formed (**Fig. 6j**). We speculate that at first, e.g., during mitotic exit, promoters of active genes interact. Subsequently, RNA polymerase II is recruited to these promoters and RNA is generated. At speckle-associated domains the high density of active genes, the high level of their expression, and possibly the nature of the RNAs, produce a glue of RNA around the locus that spreads beyond the genes, leading to stabilized chromatin interactions throughout the speckle associated domains. Finally, these domains become associated with the non-chromatin nuclear speckle body, possibly via specific RNAs such as Alu RNAs, and through contacts mediated by specific DNA binding proteins. Given that some transcription factors can mediate locus-speckle interactions independent of transcription^65^, some locus-speckle contact may already occur prior to gene activation.

Our study shows a structural and functional connection between chromatin (sub-) compartments detected by Hi-C and non-chromatin nuclear bodies such as speckles. Although the biophysical process of phase separation likely contributes to form most if not all types of chromatin compartments, it has been difficult to identify the molecular mechanisms and components driving chromatin interactions that underlie compartmentalization. Different molecules likely drive formation of different compartment types. This is illustrated by our results where the speckle sub-compartment is stabilized by RNA, while the “interior active 2/3” and lamin-associated sub-compartments are not. Future studies can reveal which RNAs are involved in speckle sub-compartment formation, and the biophysical and molecular mechanisms by which they stabilize chromatin interactions. The ability to detect chromatin interaction stability with LC-Hi-C, together with targeted perturbations can lead to the identification of molecules driving formation of other sub-compartments.

## Methods

### Experimental methods

#### Cell Culture and treatments

K562 cells (ATCC, CLL-243) were cultured at 37 °C (5% CO2 and 90% humidity) in RPMI 1640 medium with high glucose (Gibco, 11875-093), supplemented with 13% (v/v) heat-inactivated fetal bovine serum (FBS; Gibco, 16000-069), 1% penicillin-streptomycin (Gibco, 15140-122), and 1% L-glutamine (Gibco, 25030-081). All cell lines were routinely tested for mycoplasma contamination using the MycoAlert™ PLUS Mycoplasma Detection Kit (Lonza, LT07-518) and were consistently confirmed to be free of contamination. Asynchronous K562 cells at densities of 0.4-0.5M cells/ml were treated with both 25 uM triptolide (Sigma, 645900-1MG) and 200 uM 5,6-Dichlorobenzimidazole 1-β-D-ribofuranoside (DRB; Sigma, D1916-50MG) for 4 hours.

Anti-sense morpholinos (U1 and control; seeSupplementary Table 3) for oligo sequences) were obtained from GeneTools and transfected into K562 cells using the Neon transfection system (Thermofisher, MPK10096). The U1 AMO was originally designed and successfully used to inhibit splicing^67^. Briefly, cells were washed once with DPBS (Thermofisher, 14190-250) and resuspended in electroporation buffer (R buffer) at a density of 5M cells/ml. Cells were electroporated 100ul at a time in 100ul Neon tips and immediately transferred to fresh media without antibiotics. Cells were cultured for 4 hours in media without antibiotics at a final density of 0.5M cells/ml. Heat-sock experiments were carried out by adding small volumes of dense cell suspensions to plates/flasks pre-heated to 42C in a cell culture incubator. The final densities of cultures were 0.8 cells/ml. Cells were treated with 1 mM of either dTAGV-1 (Tocris, 6914) or dTAGV-1-NEG (Tocris, 6915) for 6h with or without an additional 4hr incubation with either DMSO or 25 uM triptolide/200 uM DRB.

#### Plasmid construction and electroporation

Single gRNAs were designed to target nuclear speckle-associated regions that either destabilize [Chr19: 49,465,034 - 49,475,273 with three targeting sequences - C19-d01 (5’-GAGATGGACAGnGG-3’), C19-d05 (5’-CCnCCAGAGGCTCAT-3’), and C19-d06 (5’-CCnCAGAGGCTCATG-3’)] or retain stability [Chr19: 44,800,810 - 44,801,836 with two targeting sequences - (C19-G03 (5’-CCnGGAGTCCATACT-3’) and C19-22 (5’-CCnTACTGCAGCCC-3’)] after transcription block as determined by LC-Hi-C (**Supplementary Table 1**). All targeting sequences were first cloned into the single guide RNA expression vector, pPUR-hU6-sgRNA-Sirius-8XMS2 (Addgene #121942), a pLKO.1 lentiviral-based system described in prior work^48^. We then cloned multiple proximity-targeting sgRNAs into one CRISPRainbow DONOR1 vector^68,69^ (addgene #75398) (**Supplementary Fig. 2**) to enhance the signal-to-noise ratio of the low-repetitive loci in K562 cells. The forward and reverse primers used for the PCR amplification of the C19-S and C19-DS guide fragments are listed in **Supplementary Table 4**. The sgRNAs were designed using CRISPRbar (http://genome.ucf.edu/CRISPRbar/)^48^. The assembled CRISPRainbow-DONOR1--sgRNA plasmids and dCas9 expression vector pHAGE-TO-dCas9-P2A-HSA (Addgene #121936) were delivered into cells via electroporation using Neon™ Electroporation System (Invitrogen, MPK10096), following the manufacturer’s protocol.

#### Lentiviral packaging

Lentiviral particles were produced in HEK293T cells as previously described^48^. Briefly, approximately 0.5 x 10^6^ HEK293T cells per well were seeded into 6-well plates 24 hours prior to transfection. Cells were co-transfected with 0.5 µg of pCMV-dR8.2 dvpr (Addgene #8455), 0.3 µg of pCMV-VSV-G (Addgene #8454), and 1.5 µg of pHAGE-EFS-MCP-tdTomato using TransIT transfection reagent (Mirus), following the manufacturer’s instructions. Viral supernatants were harvested 48 hours post-transfection, filtered through a 0.45 µm filter, and either used immediately or snap-frozen and stored at –80 °C.

#### Generating a stable cell line carrying MCP-tdTomato

We established a K562 cell line, K562-MCP-tdTomato, for tracking genomic locus dynamics. We transduced lentiviral particles carrying the MS2 capsid protein-expressing vector pHAGE-EFS-MCP-tdTomato into K562 cells by spinfection in 6-well plates. Approximately 2 × 10 cells were incubated with 1 mL of lentiviral supernatant and centrifuged at 1,200 × g for 30 minutes. Selection was carried out by identifying red-orange fluorescence from tdTomato (**Supplementary Fig. 3**). Fluorescence-activated cell sorting (FACS) was conducted using a BD FACSAria Fusion Flow Cytometer equipped with 405 nm, 488 nm, 561 nm, and 637 nm excitation lasers. MCP-tdTomato was excited with the 561 nm laser and detected using a 575/26 nm emission filter (the PE-YG laser channel).

#### Mounting K562 cells on coated imaging dishes

To prepare the live cell imaging dish, 250 μL of Poly-L-Lysine solution (Sigma-Aldrich, P6282-5MG; stock concentration 0.1 mg/mL) was evenly spread to the center of a 35 mm glass-bottom dish (MatTek, P35G-1.5-14-C). Afterward, 200 μL of the solution was carefully removed, leaving approximately 50 μL evenly distributed at the center. The dish was incubated for 5 minutes at room temperature, after which the remaining solution was removed using vacuum aspiration. The surface was then thoroughly rinsed with sterile tissue-culture grade water, and all residual liquid was aspirated. The coated dish was air-dried for at least 2 hours in a sterile hood before cell seeding. For imaging, 250 μL of K562 cells at a concentration of 1×10 to 6×10 cells/mL were gently added to the center of the coated dish. Cells were incubated for at least 15 minutes at 37 °C in a 5% CO₂ incubator prior to imaging.

#### Live cell imaging

We performed imaging using an Olympus IX83 inverted microscope equipped with three EMCCD cameras (Andor iXon 897) and four laser lines (405 nm, 488 nm, 561 nm, and 647 nm). The system included a 1.6X magnification adapter and a 60× apochromatic oil immersion objective lens (NA 1.5), corrected for coverslip thickness and temperature, yielding a total magnification of 96X. The microscope stage was enclosed in an incubation chamber maintained at 37 °C with controlled CO₂ and humidity. Image acquisition was carried out using CellSens software. Images were cropped to display individual nuclei. Contrast and intensity thresholds were adjusted to optimize visualization of nuclear signals over background fluorescence. Post-acquisition image processing and data analysis were performed using Fiji^70^ and Mathematica (Wolfram v12.3.1). **Fig. 3g** and **Extended Data Fig. 3g** were generated using OriginLab (2024).

#### Immunofluorescence microscopy

K562 cells were seeded onto poly-L-lysine (Sigma, P1274-25MG) coated coverslip in 6-well plates using cultures with densities between 0.4-0.8M cells/ml and allowed to adhere to the coverslips for one hour in a cell culture incubator. Coverslips were fixed in 4% paraformaldehyde (Fisher, 15710) in 1xPBS for 10 minutes followed by five 1xPBS (Thermofisher, 70013-073) washes. When immunofluorescence staining was combined with RNA-FISH, coverslips were first rinsed in CSK buffer (10.35% w/v sucrose, 10mM PIPES pH7.8, 100mM NaCl, 3mM NaCl), then pre-permeabilized in CSK + 0.5% v/v Triton X-100 + 5% v/v vanadyl ribonucleoside complex (VRC; NEB, S1402S) for 3-5 minutes before fixing cells in 4% paraformaldehyde in 1xPBS for 10minutes. Coverslips were then rinsed once in 70% ethanol then stored in 6-well plates containing 70% ethanol. For immunofluorescence on isolated nuclei, mock-or DpnII-digested nuclei were fixed with paraformaldehyde at a final concentration of 4% for 20 min. Nuclei were carefully layered onto 30% sucrose in nuclei buffer [10 mM PIPES pH 7.4, 10 mM KCl, 2 mM MgCl2, 1 mM DTT and 1X Protease inhibitor (Thermofisher; 78429)] in a microplate well containing a poly-L-lysine-coated coverslip and spun down for 15min at 2500xg at 4 °C. Coverslips were washed five times with 1xPBS then stored at 4 °C before immunofluorescence staining. Cells were then permeabilized in 0.5% Triton X-100 (Sigma, 93443-500mL) for 10 minutes before blocking with 3% BSA (Sigma, A7906-100G) in 1xPBS for 1 hour. Cells were then labeled with primary antibody (**Supplementary Table 5**) diluted in 3% BSA in 1xPBS for 1-3 hours as per manufacturer recommendations. Coverslips were washed three times with 1xPBS before staining with fluorescently labelled secondary antibody diluted in 3% BSA in 1xPBS for 1 hour. Coverslips were washed three times with 1xPBS before staining with 1 ug/ml DAPI (Thermofisher, EN62248), washing three times with 1xPBS and mounting with Vectashield mounting media (Vector Labs, H-1000-10).

When immunofluorescence staining was combined with RNA-FISH, immuno staining was done as above with a few exceptions. In this case we employed the use of Ultrapure BSA (Thermofisher, AM2618) and included 1U/ul RNasIN (Promega, N2515)) when blocking and antibody staining. A 1xPBS wash followed by a wash with 1xPBS+0.1% Triton X-100 and another 1xPBS wash was done after staining for primary and secondary antibodies. After the secondary antibody washes, cells were fixed again in 4% paraformaldehyde in 1xPBS for 10 minutes, washed twice in 1xPBS before continuing to RNA-FISH. When immunofluorescence staining was combined with DNA-FISH, coverslips were fixed using 2% paraformaldehyde in 1xPBS. Immunofluorescence staining was done as above with the exception that, after secondary antibody staining, coverslips were subjected to a 1xPBS wash followed by a wash with 1xPBS+0.1% Triton X-100 and another 1xPBS wash. After the washes, cells were fixed again in 4% paraformaldehyde in 1xPBS for 10 minutes, washed twice in 1xPBS before continuing to DNA-FISH.

#### DNA FISH

Coverslips carried over from Immunofluorescence were placed face-down onto a drop of 0.1ug/ul RNase A diluted in 1xPBS in a 6-well plate lined with parafilm and incubated at 37 °C for 1 hour. Coverslips were then rinsed three times with 1xPBS followed by permeabilization in 0.7% Triton X-100/0.1N HCl for 10 minutes on ice. Coverslips were washed three times in 1xPBS then denatured in 1.9N HCl for 30 minutes on the bench. Coverslips were washed again three times in 1xPBS. Coverslips were placed face-down onto a drop of probe mix (2ul of 10 pmol/μl probe [designed and synthesized by Daicel Arbor biosciences (**Supplementary Table 6**)] in hybridization buffer [40ul of 50x Denhardt’s media {1% ficoll (Thermo Scientific, B22095.09), 1% polyvinylpyrrolidone (Sigma, P2307-100G), 1% BSA}, 160ul of 25% Dextran sulfate (Fisher, BP1585-100) and 200ul formamide (Sigma, S4117)]), sealed in parafilm, placed in a humid chamber and incubated in a 37 °C oven overnight. Coverslips were then washed for 30 minutes in 0.5x SSC + 0.1% Tween-20 pre-warmed at 37 °C. Coverslips were then washed sequentially with 2xSSC + 0.1% Tween-20, 1xSSC + 0.1% Tween-20 then 0.5xSSC + 0.1% Tween-20 for 30 minutes each at room temperature. Finally, coverslips were washed once in 1xPBS before staining with 1 ug/ml DAPI (Thermofisher, EN62248), washing three times with 1xPBS and mounting with Vectasheild mounting media (Vector Labs, H-1000-10).

#### RNA FISH

Coverslips carried over from Immunofluorescence staining were dehydrated in ice-cold 100% EtOH for 10 minutes then air dried until visibly dry. Coverslips were rinsed with wash buffer [10% formamide (Sigma, S4117), 2x SSC] for 10 minutes before adding 100ul of probe mix (Biosearch Technologies, SMF-2035-1), covering with parafilm and incubating in a 37 °C oven for 3 hours. Coverslips were then washed with wash buffer for 30 minutes at 37 °C. Finally, coverslips were washed twice in 1xPBS before staining with 1 ug/ml DAPI (Thermofisher, EN62248), washing three times with 1xPBS and mounting with Vectashield mounting media (Vector Labs, H-1000-10).

#### Confocal microscopy

Imaging was performed on a Leica Stellaris 8 confocal laser scanning microscope (Leica Microsystems) using a HC PL APO CS2 100x/1.40 OIL objective. Fluorescence was excited using a 405 nm laser at 2.30% power and a white light laser tuned to 499 nm, 579 nm and 653 nm at 6.31%, 5.77% and 15.35% laser power, respectively. Emitted signals were collected sequentially using Leica detectors with the following spectral detection windows: HyD S Channel 1,: 430nm - 504nm; HyD X Channel 2: 504nm - 573nm; HyD S Channel 3: 585nm - 663nm; HyD X Channel 4: 663nm - 750nm. The confocal pinhole was set to 1.0 Airy Unit (152.7 um) for each channel. Images were acquired at a resolution of 432 × 432 pixels with a pixel dwell time of 1.575 µs and 2-line averaging applied to reduce noise [CB1.1]. For three-dimensional reconstruction, optical sections varying in total depth were taken where z-stacks had a z-step size of 0.3 µm. Computational deconvolution was performed in real time using the Leica Lightning module within LAS X, which applies an adaptive, image-content-based algorithm to enhance spatial resolution and signal-to-noise ratio beyond the classical diffraction limit. All acquisition and deconvolution parameters were controlled using Leica Application Suite X (LAS X, version 1.4.6.28433).

#### Liquid chromatin Hi-C

The Hi-C/Liquid Chromatin Hi-C (LC-Hi-C) protocol largely follows Belaghzal et al.^29,71^, respectively, with some modifications. K562 cells were harvested when undergoing exponential phase growth unless otherwise specified.

##### Nuclei digestion and processing

Nuclei were thawed on ice, 1ml of HBSS (Thermofisher, 14025134) was added, nuclei suspension was mixed gently by pipetting and samples were spun at 1,500xg for 8mins. Nuclei were washed once with 1xNEB r3.1 buffer (NEB, B6003S), counted and resuspended at a density of 6M nuclei/ml. For the samples protected by RNase inhibitor, 0.8U/ul RNasIN (Promega, N2511) was added to each solution. For RNase depletion, RNaseA (10mg/ml; Sigma, 1010969001) was added to a final concentration of 100 ug/ml and an equivalent volume of 1x NEB r3.1 was added to the control (mock-digested) samples. DpnII (50U/ul; NEB, R0543M) was added to a final concentration of 0.24U/ul for the DpnII-digested samples and an equivalent volume of 1x NEB r3.1 was added to the mock-digested samples. Aliquots of 500ul were distributed to 1.5ml tubes and incubated at 37 °C for exactly 2 or 4hrs.

Samples of the digest were taken for 1) fragment analysis (0.5M nuclei from both Mock and DpnII-digested samples), 2) DpnII-seq (0.5M nuclei from DpnII-digest only; processed in methods section “DpnII-seq”) and 3) LC-Hi-C (3M nuclei from both mock-and DpnII-digested samples). The LC-Hi-C sample was immediately fixed using 1% formaldehyde, washed, flash frozen and stored at-80 °C. DpnII-seq samples are processed in the methods section “DpnII-seq”. DNA was extracted from the fragment analyzer samples using either a single phenol:chloroform extraction, methods which are described below.

##### Hi-C protocol

Hi-C was performed as described previously by Belaghzal et al.^71^ with minor modifications. In brief, flash-frozen cross-linked mock-and DpnII-digested nuclei were digested with DpnII at 37 °C overnight. DNA overhangs generated by DpnII-digestion were filled with biotin-14-dATP at 23 °C for 4 hrs then ligated with T4 DNA ligase at 16 °C for 4 hrs. DNA was then treated with proteinase K at 65 °C overnight to remove cross-linked proteins. Ligation products were purified using either single phenol chloroform extraction followed by ethanol precipitation and elution, or using Ampure beads without size selection. The library was fragmented by sonication to an average size of 200 bp and size-selected to fragments of 100–350 bp. We then performed end repair and dA-tailing and selectively purified biotin-tagged DNA using streptavidin beads. Illumina PE or TruSeq adapters were added to the final Hi-C ligation products, samples were amplified and the PCR primers were removed. Hi-C libraries were then paired-end sequenced using Illumina (either HiSeq 4000, NextSeq 2000, NovaSeq 6000 or NovaSeq X Plus).

#### DpnII-seq

Half a million DpnII-digested nuclei (from section “Liquid Chromatin Hi-C: Nuclei digestion and processing) were transferred to a 1.7ml tube containing 10ul of 0.5M EDTA and transferred to 65 °C for 20min, before adding 50ul of 10mg/ml Proteinase K and incubating at 65 °C 3 hours to overnight. DNA was extracted using either a Phenol:chloroform extraction followed by ethanol precipitation and column washes or using HB-Ampure mixture with a final elution in 85ul of TLE. To each DpnII-seq sample, 15ul of Fill-in mastermix was added which contains 0.5ul of ultrapure water 1.5ul 10X NEBuffer 3.1 (NEB, B7203S), 0.4ul of 10mM dCTP (Invitrogen, 56173), 0.4ul of 10mM dGTP (Invitrogen, 56174), 0.4ul of 10mM dTTP (Invitrogen, 56175), 9.4ul of 0.4mM biotin-14-dATP (LifeTech, 19524016) and 2.5ul of Klenow DNA Polymerase (NEB, M0210L). Samples were incubated for 1.5 hours at 23 °C, before adding 1ul of 10mg/ml RNase A and incubating at 37 °C for at least 30 minutes. Samples were brought up to 132ul with ultrapure water for agarose gel QC, sonication, size selection and End-repair^29^. However, instead of proceeding with pull-down samples were purified using a 2x Ampure mixture to sample ratio and eluted in 41ul of TLE. A-tailing was done followed by another 2x Ampure to sample cleanup eluting DNA in 32ul of ultrapure water. To the eluted sample, 8ul of 5x ligation buffer (Invitrogen, Y90001) and 10ul of ligation master mix was added to each sample. DpnII-seq samples were cleaned up using a 2x Ampure to sample ratio and eluted in 90ul of ultrapure water. To each library, 10ul of 10x NEB CutSmart buffer was added along with 3ul of USER enzyme and samples were incubated to 1 hour at 37 °C. DpnII-seq samples were then subject to biotin pulldown as described in the LC-Hi-C methods but final washes were done in 1x NEB r2.1 buffer instead of TLE. Illumina PE or TruSeq adapters were added, samples were amplified and the PCR primers were removed. Libraries were then paired-end sequenced using Illumina (either HiSeq 4000, NextSeq 2000, NovaSeq 6000 or NovaSeq X Plus).

##### Fragment size analysis

DNA is extracted from the aliquot of pre-digested nuclei taken during the LC-Hi-C protocol using a single phenol:chloroform extraction followed by a 30 minute incubation at 37 °C in the presence of 50 ug/ml RNase A. DNA concentration is then quantified using Qubit (Thermofisher, Q33266), samples appropriately diluted and run on an Agilent Fragment Analyzer using the HS Large Fragment 50kb kit (Agilent, DNF-464-0500).

#### SLAM-seq

Cells were lysed and resuspended in Trizol reagent (Thermofisher, 15596018). External RNA Controls Consortium (ERCC) spike-ins with and without 4sU (at a 10% 4sU:U concentration) were added before extraction to each sample at a concentration of 2.4ng per million cells. Total RNA was extracted using the TRIzol– chloroform method, followed by isopropanol precipitation and 75% ethanol washes.

RNA integrity was assessed using an Agilent TapeStation, and samples with RNA Integrity Number (RIN) ≥ 9 were used. SLAM-seq was done following the published protocol^42^. Briefly, RNA was alkylated with iodoacetamide (IAA) 10mM in a mix containing NaPO4 pH 8 50 mM and DMSO 50%. The mix was incubated at 50 °C for 15min and quenched by the addition of Dithiothreitol (DTT) to a concentration of 20mM followed by a cleanup using the RNA Clean & Concentrator-5 columns (Zymo Research, R1013) following the manufacturer’s protocol. Alkylated RNA was depleted from rRNA using an RNAseH-based RNA depletion using custom probes targeted to mammalian rRNA sequences^72^ and prepared for sequencing using the NEBNext® Ultra™ II Directional RNA Library Prep Kit (Illumina, E7760L). Samples were sequenced on an Illumina NextSeq2000 at a minimum sequencing depth of 30M reads per sample and a length of 100 paired-end.

### Genomic Analysis

#### Public data and annotations

Publicly available SPIN annotations for K562 (Wang, 2021; https://github.com/ma-compbio/SPIN) were used throughout this paper. Various publicly available RNA-seq, CHip-seq, TSA-seq and DamID data were also used and are tabulated in **Supplementary Table 7**. Candidate cis-regulatory elements (cCREs) for K562 were obtained from the ENCODE Registry of cCREs (Registry V4, hg38) using the SCREEN interface (screen.wenglab.org). Elements were classified into PLS, pELS, dELS, CA-only, CA-H3K4me3, CA-CTCF and CA-TF categories as defined by the registry^55,73^.

#### Hi-C data pre-processing

Fastq files were mapped to hg38 using the distiller-nf pipeline (https://github.com/open2c/distiller-nf/), as in Akgol Oksuz et al.^74^. Briefly, read pairs were mapped independently to hg38 using bwa mem^75^ and then parsed into read pairs using pairtools^76^. Read pairs were subsequently aggregated into cooler-format binned contact matrices at resolutions of 1, 2, 5, 25, 50,, 100, 150, 250, 500 and 1000 kb each normalized using iterative correction^77^. Contacts within 1kb distance were discarded.

#### Hi-C analysis

Hi-C analyses were scripted in Python (>= v3.7.1) mainly using the cooltools (>= v0.5.1) package within Jupyter notebooks^78^. Default parameters for functions were used unless otherwise specified. Here, we briefly describe the use of several functions in these packages used in our analysis.

##### Calculate ‘expected’ contact frequency by distance

The frequency of interactions as a function of genomic distance (*P(s)*) was calculated using balanced Hi-C data binned at 1kb using the ‘expected_cis’ function. Scaling was performed on chromosome arms to generate average genome-wide scaling. For several analyses and visualizations (including the generation of log2(observed/expected) heatmaps and saddle plots), Hi-C matrices were normalized for this expected decay of interaction frequency with genomic distance.

##### Eigenvectors and compartments

Eigenvector analysis was performed on whole chromosomes using the ‘eigs_cis’ function at 50, 150, 250 and 1000kb where gene density was used to phase eigenvectors. The EV1 vector most often corresponded to Hi-C compartments and was saved as a text file for downstream analysis and visualization.

##### Pairwise-class averaging via saddle plots

We evaluated how different classes of regions interact with each other using saddle plots generated from the ‘saddle’ function. Classes of genomic loci (e.g. compartment status or SPIN state) are first assigned on a per bin level and passed as the ‘track’ parameter to the function. The average level of interactions for each possible combination of classes is then calculated from the observed/expected contact frequency heatmaps. Pairwise class average log2(observed/expected) interaction matrix is then plotted.

The assignment of bins to specific classes depends on the class type. Continuous variables such as ‘EV1’ were digitized into 38 quantiles after excluding 2.5% of the data (i.e. extreme values) at either end of the range. On the other hand, ‘SPIN state’ assignments were used directly by classifying 50kb bins according to publicly available annotations.

We estimate the strength of A and B compartments as enrichment of homotypic (AA or BB) over heterotypic interactions [e.g. AA / ((AB + BA) / 2)]. Here, each group (AA, BB, BA, AB) is an average of observed/expected interactions between the highest or lowest 20% EV1 quantiles for a given group. For example, the AA group is an average observed/expected interaction frequency between the highest 20% of interacting EV1 quantiles, whereas the the AB (or BA) is an average of the observed/expected interactions between the highest 20% of EV1 quantiles (A compartment) interacting with the lowest 20% of EV1 quantiles (B compartment).

##### Average feature pileups

We explored interactions between candidate promoters from our cCRE annotations. The ‘pileup’ function was used to extract +/-100kb obs/exp Hi-C map snippets centered on each promoter (anchor) pair to generate a stack that is then collapsed into a mean pileup matrix. A central dot indicates enriched contacts (loops) between promoters. Loop strength is calculated by taking the log ratio of the center pixel over the average of the pixels in each of the four corners [i.e. log(center/((UR + UL + LR + LL)/4)].

#### DpnII-seq analysis

DpnII-seq libraries were analyzed using https://github.com/dlafont15/DpnII-seq. Briefly, sequenced reads were mapped to hg38 using the bowtie read aligner^79^ and reads mapping to multiple regions of the genome were removed. In order to remove artificial biases, we filtered out paired-end reads from fragments where cut-sites were more than three nucleotides away from the start or end of the paired-end reads. Filtered reads were then binned at a resolution of choice. To account for copy number variation due to karyotype and rearrangements in the K562 cell line, we leveraged publicly available copy number data from the CatalOgue of Somatic Mutations In Cancer (COSMIC) database to assign a copy number state to each genomic bin. Read coverage files were corrected to a genome-wide diploid state using the copy number assignments and dividing coverage by an appropriate correction value (i.e. diploid = 1, triploid = 1.5, tetraploid =2, etc). Final copy number corrected coverage files were used for all downstream analyses.

#### Liquid Chromatin Hi-C analysis

The analysis of LC-Hi-C in the is paper largely builds from the work done in Belaghzal et al.^29^ and was written in R and python as both executables and scripts to be run R Studio and bash, respectively. All scripts can be found at “https://github.com/dekkerlab/speckle_RNA_dynamics_paper/LCHiC/” and incorporate functions from ‘https://github.com/tborrman/liquid-chromatin-Hi-C/tree/master/src/scripts’.

##### Calculate Loss of Structure

The Loss of structure (LOS) metric quantifies the loss of interactions at the diagonal after DpnII pre-digestion at every genomic bin^29^. This is accomplished by first using ‘cool2RangeCisPercent.py’ and calculating the percent of interactions that occur within a 2Mb window along the diagonal then dividing that by the total number of interaction (percent cis) within that bin. This collapses the 2D Hi-C data to a single dimension. The ‘LOS.R’ tool then calculates LOS by first computing the difference between mock-and DpnII-digested cis percent values genome-wide then dividing by the Mock-digested percent cis value for each bin:

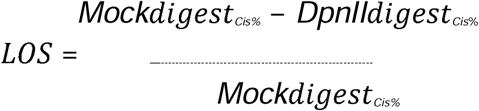

##### Calculate Loss of Contacts

The Loss of contacts (LOC) metric quantifies the loss of interactions within the diagonals of mock-digested heatmaps before and after a given treatment. This is accomplished by first using ‘cool2RangeCisPercent.py’ to calculate percent cis for mock-digested controls for a given treatment. The LOS.R script then calculates LOC by calculating the difference between cis percent values for mock-digested control and treatment then dividing by the cis percent values for the control mock-digest:

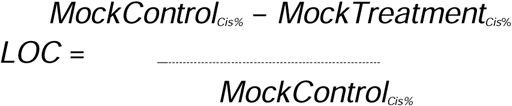

##### Calculate LOS residuals

The DpnII-seq sample is used to normalize our LOS metric. This is done by first plotting LOS against LOS, calculating a moving average and determining the residuals between the fit and the data point. These operations are done interactively using the ‘make_residual_table.R’ script.

#### Bootstrap analysis

To assess whether the difference in LOS residual values between control and treatment differed from zero within each SPIN state, we performed a bootstrap analysis. For each difference comparison, genomic bins were grouped by SPIN state and bins with missing values were excluded. To account for different group sizes, each SPIN state was resampled at the number of 150kb bins of the smallest SPIN group. Within each SPIN group we drew 10,000 samples of size equal to the number of bins representing each SPIN group. Values were drawn with replacement and mean LOS difference for each sample was computed yielding a bootstrap distribution of mean difference per state. Two-sided p-values were computed from the proportion of bootstrap means falling on the opposite side of zero from the observed mean (doubled). p-values were corrected for multiple testing across SPIN states using the Benjamini–Hochberg method.

#### Stackup and averaging features

One dimensional stacks were generated using ‘bbi.stackup’. Candidate cis-regulatory elements (cCREs) for K562 were obtained from the Search Candidate cis-Regulatory Elements by ENCODE (SCREEN) database. We defined 10kb flanks around each cCRE midpoint to generate 20kb regions which were divided into 100 bins (200bp bins). For each bin in each region, mean LOS signal (LOS data binned at 5kb) weighted by the extent to which each value covers the bin was calculated, returning a matrix where rows correspond to individual 20kb cCRE regions and the columns are the 200 positional bins. This was done separately for each cCRE class. Bins that contain all missing data are removed and rows are sorted in descending order by the average signal calculated using 5 bins on each side of the cCRE midpoint. Aggregate profiles were generated by collapsing the rows into single average values (one value per positional bin) ignoring missing values. LOS values across cCRE classes were pooled to generate an appropriate color scale across each heatmap.

For the analysis using scaled gene bodies, dominant transcripts were first inferred by determining transcript-level abundances of total RNA-seq reads for K562 (ENCSR792OIJ). Briefly, quantification was performed with Salmon (v2.0.1) against a decoy-aware index built from Ensembl GRCh38 transcript sequences (cDNA and ncRNA), using the genome as decoy. Salmon was run in mapping-based mode, with automatic library-type detection (-l A) and sequence-specific and GC-content bias corrections enabled (--seqBias, --gcBias). Transcript-level abundances (TPM) and estimated read counts were then used to identify a dominant transcript for each gene annotation. For genes with detectable expression (TPM > 0), the isoform with the highest TPM was designated as dominant. For genes with no detectable expression, the longest annotated transcript was retained. Each dominant transcript was assigned a SPIN state by overlapping its transcription start site using 50-kb SPIN annotations.

Transcripts were subset by SPIN group (Speckle vs. Interior_Act23) and by expression (not expressed, TPM = 0; expressed, TPM > 5). Dominant transcripts were further restricted to those with genomic span (end − start) greater than 10 kb.

For each transcript, the gene body (TSS to TES) was rescaled to a fixed number of bins. These scaled regions were flanked by 10-kb upstream and downstream windows, each binned at fixed resolution. Minus-strand transcripts were reversed and rows with no signal were dropped. Transcripts were sorted by mean signal in a window centered on the TSS. Mean profiles were computed across transcripts as done above. POLR2A ChIP-seq, (ENCFF496FVA) and SLAM-seq CPM (this study) were profiled over the same sorted regions generated using LOS residuals.

#### SLAM-seq analysis

Reads were aligned to HG38 v95 with STAR, with specific settings optimized for SLAM-seq^80^, adding options --outFilterMismatchNmax 20 -- outFilterScoreMinOverLread 0.4 --o outFilterMatchNminOverLread 0.4 --alignEndsType Extend5pOfReads12. T>C conversion efficiency was checked using the 4sU-labeled ERCC spike-ins. Aligned reads were processed with grand slam^81^ to obtain counts per gene with T>C conversions for both exonic and intronic regions. New-to-total RNA ratios (NTR) were only calculated for exons. Counts for exons, introns or whole genes were used to get differential gene expression using DESeq2^82^ with size factors set to the mean number of reads for the ERCC spike-ins.

### Imaging Analysis

#### 3D volume analysis

Representative images were generated using Fiji^70^. As SON intensity varied across images of same treatment, image stacks were first normalized for intensity using a custom python workflow. Briefly, a single global normalization factor for the SON channel was computed for each stack. We then divide every voxel by this factor to preserve the relative intensity differences between z-slices. These normalized images were then saved as tif files and imported into Aivia (v13.0.0) for 3D segmentation and volume analysis using the “3D object analysis” tool. Speckle volume and speckle count per cell data are then extracted from the Aivia output and plotted.

#### Quantifying SON intensity at DNA foci

Stacks obtained from imaging of cells stained with both anti-SON antibody and DNA-FISH probes were manually curated to extract slices containing DNA foci. The final group slices were selected so as to not duplicate foci. If a single foci spanned several slices, the slices containing the highest SON signal surrounding the foci were selected. Images were further curated to maximize the number of segmented foci while removing images with segmented background signal. Imaging files containing three channels (DAPI, SON and DNA FISH) were processed using custom jupyter notebooks (see ‘FISH_quant.ipynb’). Briefly, intensity normalization factors across all slices are first calculated independently for the DAPI and DNA FISH channels. Reference values (99th percentile intensity values) for each slice are determined, then scaled to the mean of these values across all slices. Slices from the DAPI and DNA-FISH channels are then filtered and segmented using the ‘skimage’ package from scikit-image. First, a gaussian blur is applied to the DAPI channel in order to reduce noise before Otsu segmentation and filtering of small objects. The DNA-FISH signal is then percentile thresholded where 3% of the brightest signal within the nucleus (within DAPI segmentation) is segmented. A dilated mask of 5 pixels is then generated surrounding each DNA FISH foci using the ‘skimage.morphology.disk’ and ‘skimage.morphology.binary_dilation’ functions. Mean SON intensity both within (segmentation) and surrounding (dilated mask) the DNA FISH foci is calculated.

#### Analysis of genomic-locus dynamics

To eliminate the drift movement of cells, the movement of individual genomic loci was calibrated relative to the nuclear centroid. Each locus was tracked over 120 consecutive image frames with a lag time of 0.2 seconds. The mean square displacement (MSD) of lag time kΔ*t* was calculated as follows:

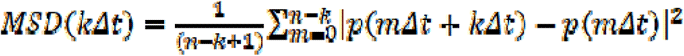

where p(t) is the position vector of a locus at time *t*, and Δ*t* is the lag time, the fixed time interval between two successive image frames. All MSD curves were fitted using the power-law equation: MSD( )=4D_app_ *( )*^α^, where D_app_ is the apparent diffusion constant and =kΔ*t*. The gyration (or trajectory) radius R_g_ of the locus trajectory was calculated as:

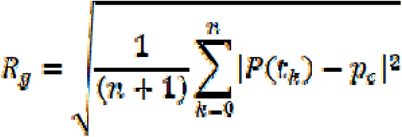

where 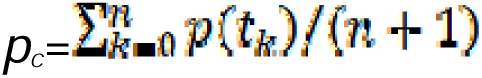 is the geometric center of the positions defining the trajectory and *t_k+1_*=*t_k_+*Δ*t*. The gyration radius R_g_ measures the size of the area covered by locus movement. The effective diffusion constant D_eff_ is calculated from MSD at the short time intervals, MSD^(s)^, within 3Δ*t* and is fitted using the equation: MSD^(s)^=4D_eff_ Δ*t*.

### Fragment Size Analysis

#### Plotting bioanalyzer traces

Since functionality for scaling and plotting fragment size distribution curves in Agilent PROsize software is limited, we employ a custom R script to calibrate and normalize the migration time over fluorescence output from Agilent Fragment Analyzer for plotting. Briefly, for each sample of a Fragment analyzer run, migration time over relative fluorescence data is extracted from ‘.raw’ files manually using PROsize software. For each Fragment Analyzer run, the migration time of each peak in the standard ladder is also obtained manually from the.raw files using the PROsize GUI. Piecewise linear functions are then calculated between each peak in the ladder to predict size (in bp) from migration time for specific migration time intervals.

The linear models generated from the ladders are then used to convert migration times into fragment sizes for each sample of that run. To compare samples between runs, ladder migration times for each peak across the ladders (of each run to be compared) are averaged to generate an averaged calibration curve. Piecewise functions are again generated between each peak of the averaged calibration ladder, but this time to convert predicted size to a common normalized time axis so we can directly plot and compare samples from different runs. Relative fluorescence is then scaled to the highest non-marker peak which is most often the maxima generated by DpnII digestion. Upper and lower bounds are then scaled to the upper and lower markers, respectively, of the Agilent sample loading buffer for HS Large Fragment kits. Scaled time curves are then plotted and the average migration time for each peak in the averaged calibration ladder are used to label and tick the x-axis with fragment size values.

## Data availability

The datasets generated in this publication have been deposited in the NCBI GEO and accession number(s) will soon be available. Published datasets are listed in Supplementary Table 7. All other data supporting the findings of this study are available from the corresponding author on reasonable request.

## Code availability

Open2C scripts and notebooks used in this study are publicly available in GitHub: https://github.com/open2c and https://github.com/dekkerlab/speckle_RNA_dynamics_paper.

## Acknowledgements

We thank the Dekker, Pai and Tu labs for helpful discussions and advice. We thank Sergey Venev for help and support for data analysis, esp. for cool2cisrange.py. We thank Chan-Wang Jerry Lio (Ohio State University) for suggestions on electroporation of K562 cells and Daniel Schoenberg (Ohio State University) for providing access to the Neon Electroporation instrument. We thank Mark Parthun and the Department of Biological Chemistry and Pharmacology at Ohio State University for the use of FACSAria Fusion Flow Cytometry. We also thank Madhoolika Bisht for her initial work during the early stages of this work. Imaging data was acquired thanks to the UMass Chan Medical School Sanderson Center for Optical Experimentation (SCOPE) on the Leica STELLARIS 8 STED confocal microscope funded by a The Umass Deep Sequencing Core was instrumental for data acquisition. We thank the Watson lab (Umass) for access to their Illumina sequencing instrument. We thank Brian Liau (Harvard University) for providing us with the K562 SON-FKBP12-F36V SRRM2-FKBP12-F36V cell line.

## Author contributions

D.L.L and J.D. conceived the project. D.L.L. and L.Y. performed genomics experiments. D.L.L performed imaging experiments. D.L.L. and with J.D. analyzed data. E.C-R. performed and with A.A.P. analyzed SLAM-seq data. Y-C.C. and N.W. performed live cell imaging experiments. Y-C.C. and L-C.T. analyzed live cell imaging data. D.L.L. and J.D wrote the manuscript with input and edits from E.C-R., Y-C.C,, L-C.T, and A.A.P.

## Extended Data Figures

**Extended Data Fig. 1:**
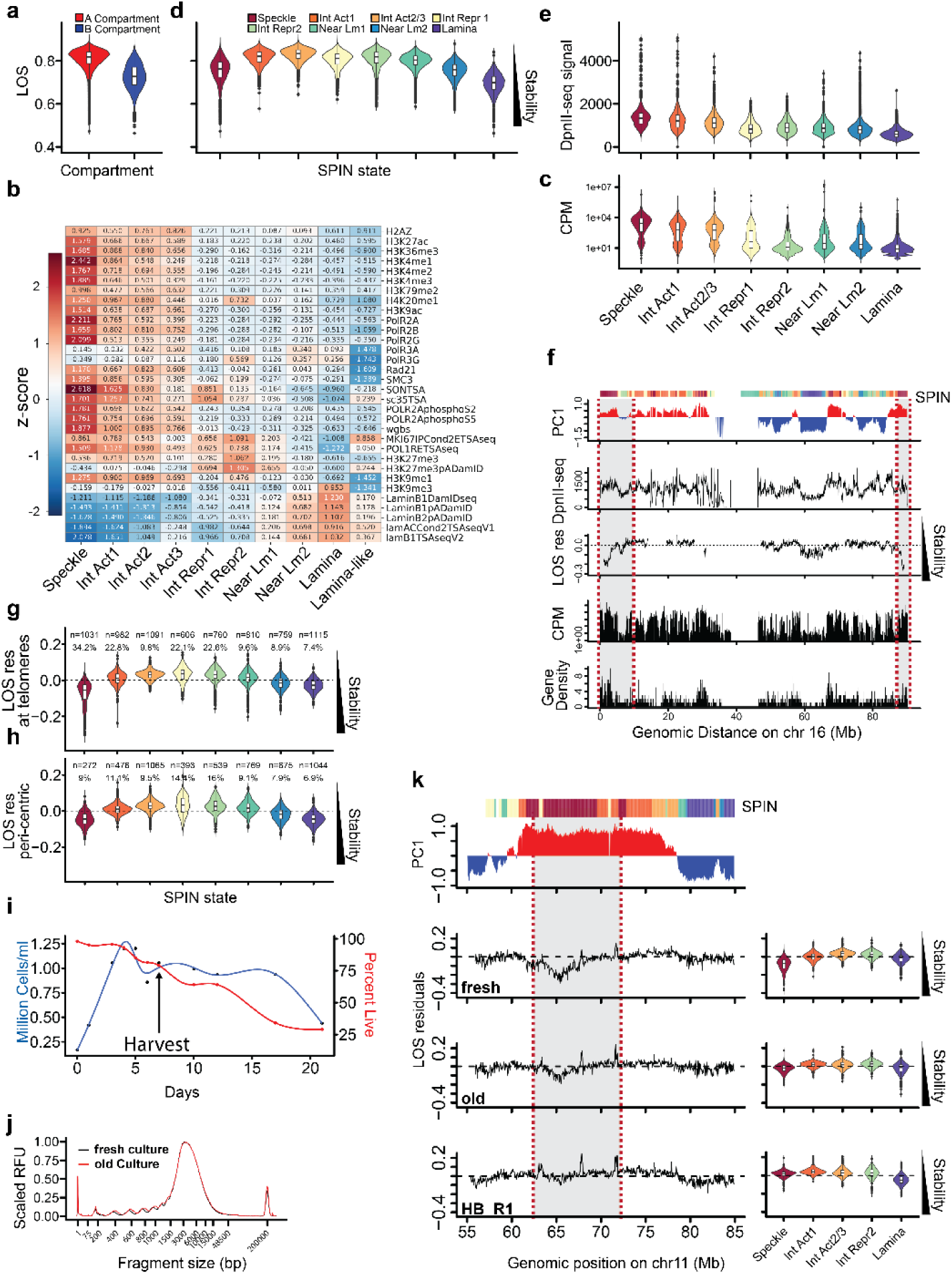
SPIN states reveal growth rate-dependent speckle-associated chromatin. **a**, LOS values grouped by compartment status (PC1). **b,** Heatmap generated using z-score normalized publicly-available data (**Supplementary Table 7**) are grouped by SPIN state. **c**, Spike-in normalized CPM values from nascent reads (SLAM-seq; see Methods) by SPIN state. **d**, LOS values grouped by SPIN state. **e**, DpnII-seq signal grouped by SPIN state. **f**, Tracks of chromosome 16 showing compartment status (PC1), SPIN state annotations in the colored bar above along with DpnII-seq signal, LOS residuals, average TPM and gene density. Speckle-associated chromatin (red dotted lines) is highlighted. LOS residuals and representation of SPIN states for bins within 10 Mb of chromosome ends (**g**) and centromeres (**h**). The number of annotated 50 kb bins from each SPIN state (n) and the percentage of annotated bins (%) are noted. **i**, Culture density and the percentage of live cells were monitored along a 21 day period without culture passage or the addition of fresh media. Harvest point for Imaging and LC-Hi-C is denoted by the arrow. **j**, Fragment size distributions generated from LC-Hi-C pre-digestion of isolated nuclei harvested from fresh/actively dividing cells (fresh), cells that were starved and unpassaged for 7 days (old). **k**, Tracks of chromosome 11 and global LOS residuals from LC-Hi-C libraries generated from Belaghzal et al (2021) (HB_R1). Speckle-associated domain is highlighted (red dotted lines). All boxplots show the median (center line), interquartile range (box), and 1.5×IQR whiskers, with outliers plotted individually. All data is binned at 50kb.

**Extended Data Fig. 2:**
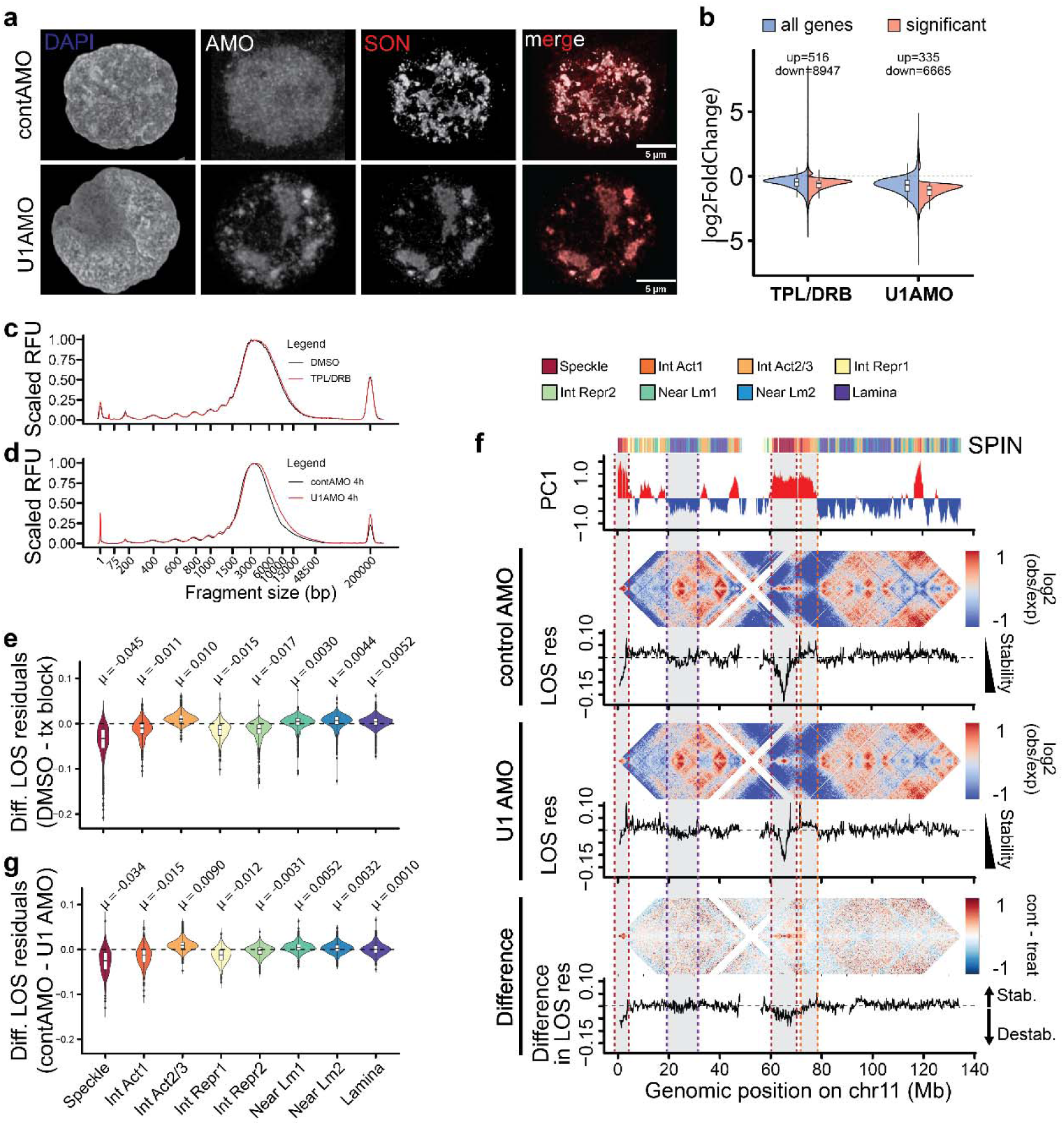
Chromatin interactions at nuclear body-associated regions differentially destabilize after transcription block and splicing inhibition. **a**, 3D projected image stacks of cells treated with either fluorescein-conjugated control AMO (cAMO) or U1AMO for 4h and labelled with an antibody against SON and DAPI. **b**, Log2(FoldChange) values after differential gene expression analysis for ‘all genes’ (protein and non-protein coding) and ‘significant’ differentially expressed genes after TPL/DRB and U1 AMO treatments using reads originating from exons. Fragment analyzer profiles of DNA fragments generated from LC-Hi-C pre-digestion of isolated nuclei harvested from cultures treated with TPL/DRB (**c**) or U1 AMO (**d**) along with their respective controls. **e**, Global difference in LOS residuals after TPL/DRB treatment for all SPIN states. **f**, log2(observed/expected) contact frequency heatmaps aligned with tracks for chromosome 11 showing SPIN states (bar above), PC1, and LOS residuals for control AMO-and U1 AMO-treated cells. The bottom heatmap and track show the difference between control (control AMO) and treated (U1 AMO) log2(observed/expected) contact frequency heatmaps and LOS residuals, respectively. **g**, Global difference in LOS residuals after U1AMO treatment (control - treatment) for all SPIN states. Bootstrap analysis generated significant p values (p < 0.0001) for comparisons of each SPIN group to 0. All boxplots show the median (center line), interquartile range (box), and 1.5×IQR whiskers, with outliers plotted individually. LOS tracks and SPIN violin plots are binned at 150kb. SPIN and PC1 tracks are binned at 50kb. Heatmaps are binned at 250 kb.

**Extended Data Fig. 3:**
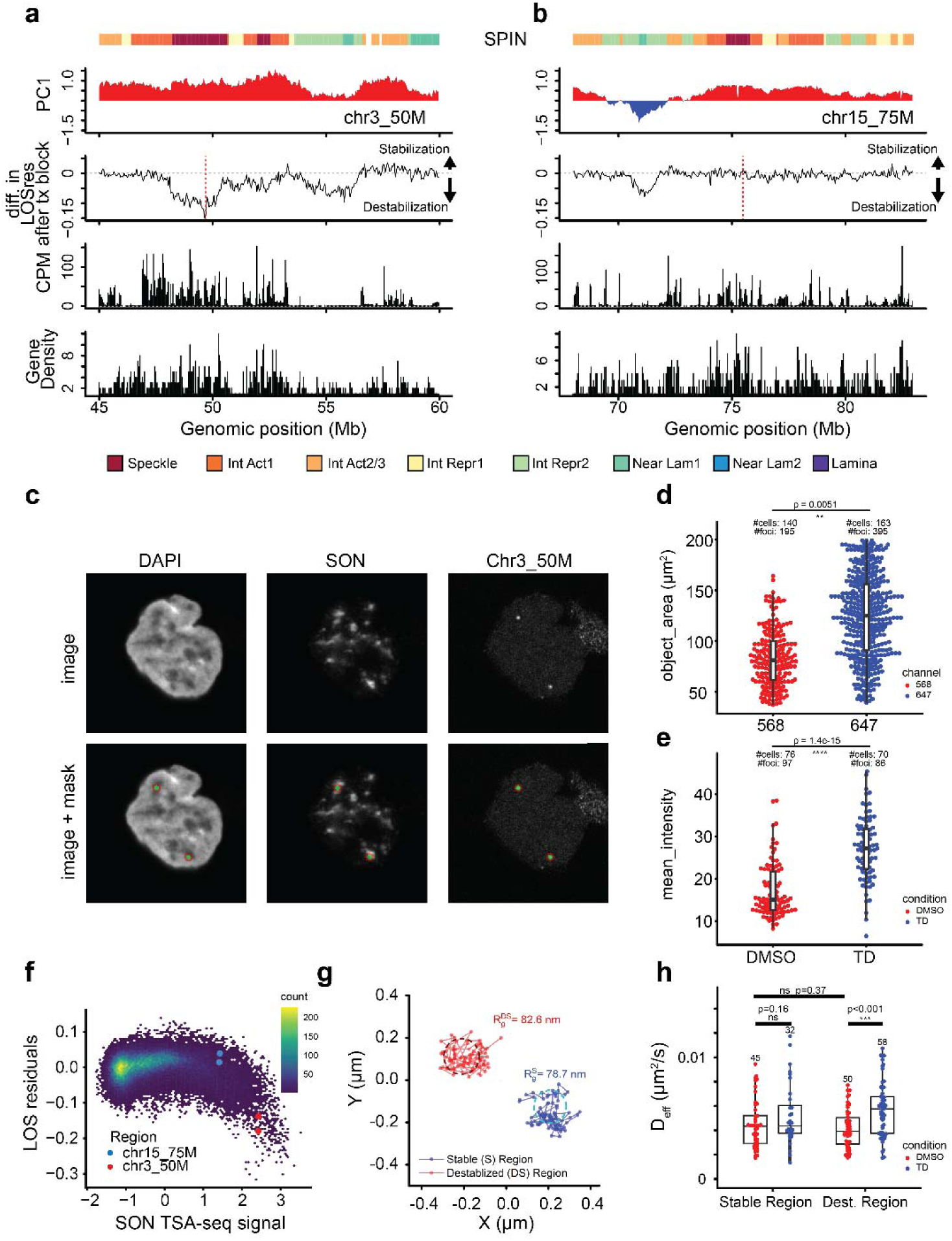
Characteristics of foci generated by FISH probes and mobility metrics. Two loci were selected for DNA FISH probe design: A 120 kb region on chromosome 3 (**a**, chr3_50M; red dotted line) that is destabilized after transcription block (TD) and a 40 kb region on chromosome 15 (**b**, chr15_75M: red dotted line) at which stability is unaffected (see **Supplementary Table 1**). SPIN state annotations, PC1, the difference in LOS residuals after transcription block (DMSO – TD), CPM for nascent SLAM-seq reads and gene density are shown for these two regions. **c,** Example segmentation of DNA-FISH foci and the surrounding 5px dilated mask. **d**, Area of DNA foci quantifications for each probe set [568 (chr3_50M) = 140 cells, 647 (chr15_75M) = 163 cells]. Kolmogorov-Smirnov (K-S) tests were performed between measurements from each group (p = 0.0051 where * p-value < 0.05, ** p-value < 0.01, *** p-value < 0.001). **e**, Image mean intensity for the SON channel in control (DMSO) and treatment (TPL/DRB) samples (KS test p = 1.4e-15; DMSO = 76 cells, TD = 70 cells). **f**, Scatter plot of LOS residual values by SON TSA-seq signal genome-wide for K562 cells. The TSA-seq values for chr19-S (blue) and chr19_DS (red) loci are denoted. **g**, Trajectories of chr19-S and chr19-DS loci and resulting D_eff_ values (**h**) after DMSO and TPL/DRB treatments (N_chr19-Stable-DMSO_=45, N_chr19-Stable-TX_=32, N_chr19-Destable-DMSO_=50, N_chr19-Destable-TX_=58). All boxplots show the median (center line), interquartile range (box), and 1.5×IQR whiskers, with outliers plotted individually. All genomic data is binned at 50 kb.

**Extended Data Fig. 4:**
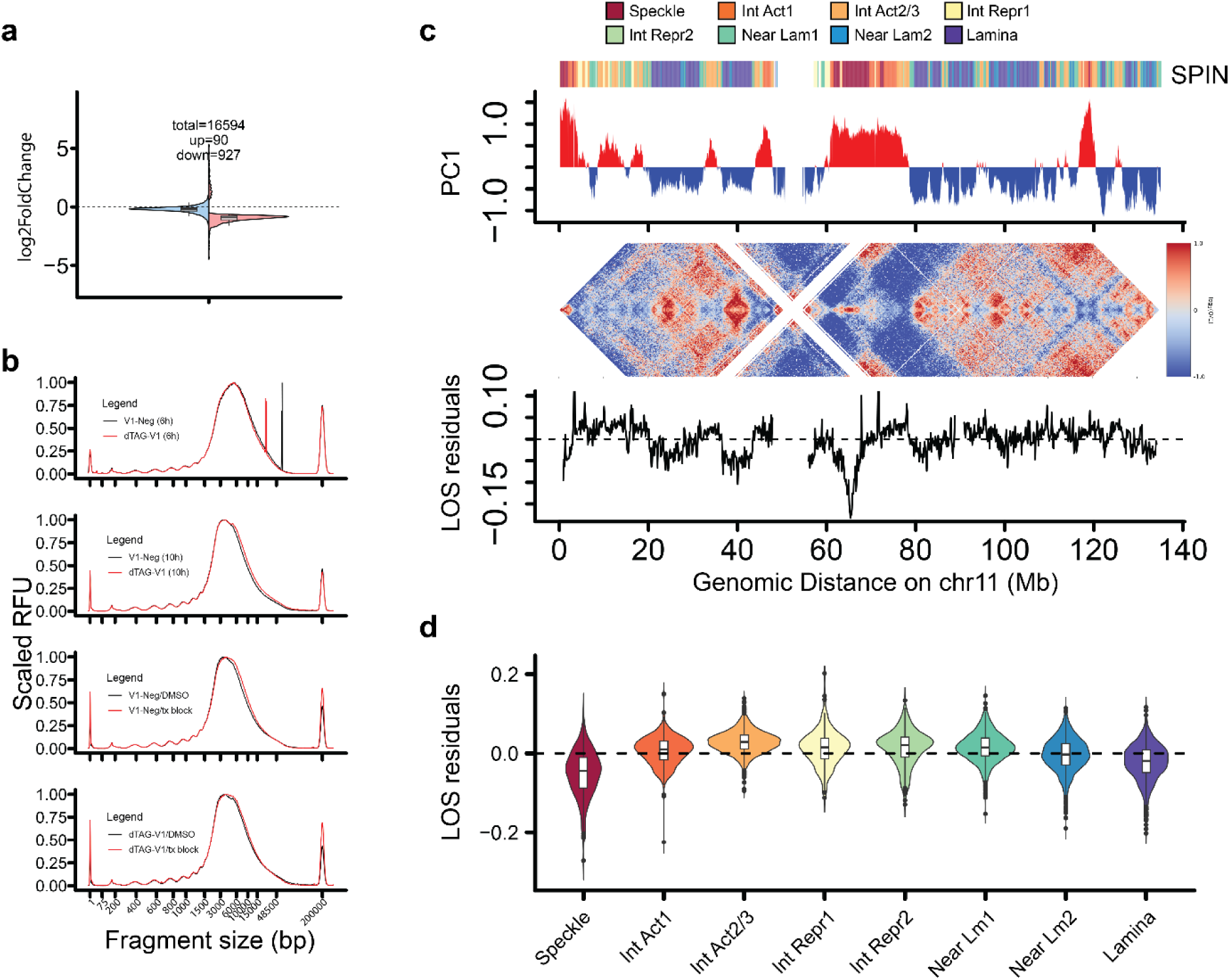
Chromatin interaction stability is unaffected by speckle depletion/dissolution. **a**, Log2(FoldChange) values after differential gene expression analysis for ‘all genes’ (protein and non-protein coding) and ‘significant’ differentially expressed genes after 6h SON and SRRM2 depletion for exonic reads. **b**, Fragment analyzer profiles of DNA fragments resulting from LC-Hi-C in situ pre-digestion of unfixed nuclei isolated after 6h SON/SRRM2 depletion [dTAG-V1(6h)],10h SON/SRRM2 depletion [dTAG-V1(10h)], 4h TPL/DRB treatment (V1-Neg/tx block) and 6h SON/SRRM2 depletion followed by 4h TPL/DRB treatment (dTAG-V1/tx block) with respective controls. **c**, Tracks of chromosome 11 with PC1, SPIN annotations and LOS residual tracks generated from control K562 cells with endogenous SON-and SRRM2-FKBP12^F36V^ edits. **d**, Global LOS residuals by SPIN state generated from control K562 cells with endogenous SON-and SRRM2-FKBP12^F36V^ edits. Bootstrap analysis generated significant p values (p < 0.0001) for comparisons of each SPIN group to 0. All boxplots show the median (center line), interquartile range (box), and 1.5×IQR whiskers, with outliers plotted individually. LOS tracks and SPIN violin plots are binned at 150kb. SPIN and PC1 tracks are binned at 50kb. Heatmaps are binned at 250 kb.

**Extended Data Fig. 5:**
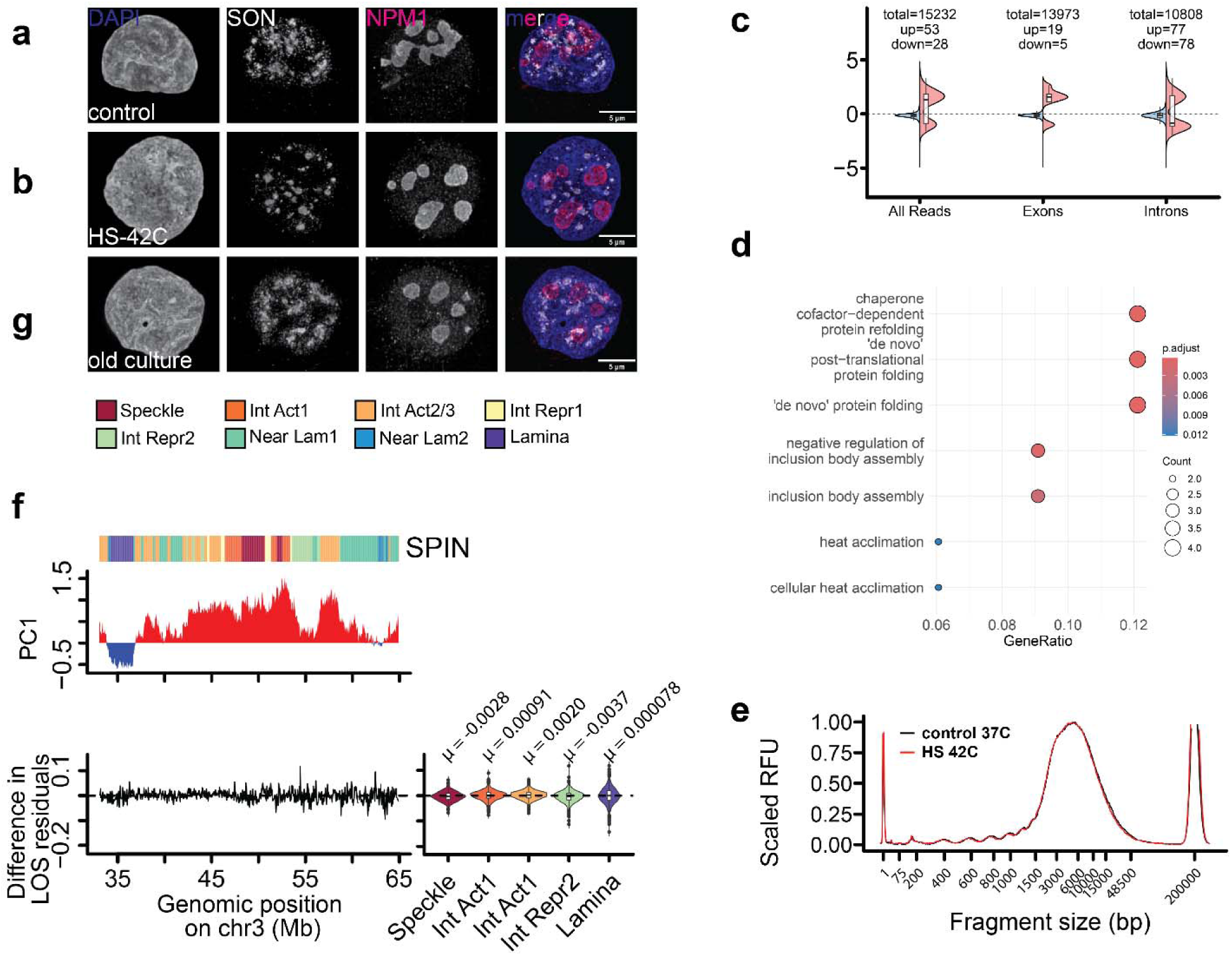
Nuclear body integrity does not always correlate with changes in chromatin interaction stability. 3D projections from image stacks of control cells (**a**) and cells transferred to 42C for 80 minutes (**b**) before labelling with antibodies against both SON and nucleophosmin (NPM1). **c**, Log2(FoldChange) values after differential gene expression analysis for ‘all genes’ (protein and non-protein coding) and ‘significant’ differentially expressed genes after 80 min heat-shock using all reads, reads originating from exons and reads originating from introns. **d**, GO-term enrichment analysis of genes upregulated after heat-shock. **e**, Fragment analyzer profiles of DNA fragments resulting from LC-Hi-C in situ pre-digestion of unfixed nuclei isolated after 80min heat-shock with respective control. **f**, Tracks of chromosome 3 (left) and global (right) difference in LOS residuals for heat-shock with respective control (control – treatment). **g**, 3D projections from image stacks of cells grown without passage or media change for 7 days before labelling with antibodies against both SON and nucleophosmin (NPM1) and staining with DAPI. All boxplots show the median (center line), interquartile range (box), and 1.5×IQR whiskers, with outliers plotted individually. Tracks are binned at 50 kb.

**Extended Data Fig. 6:**
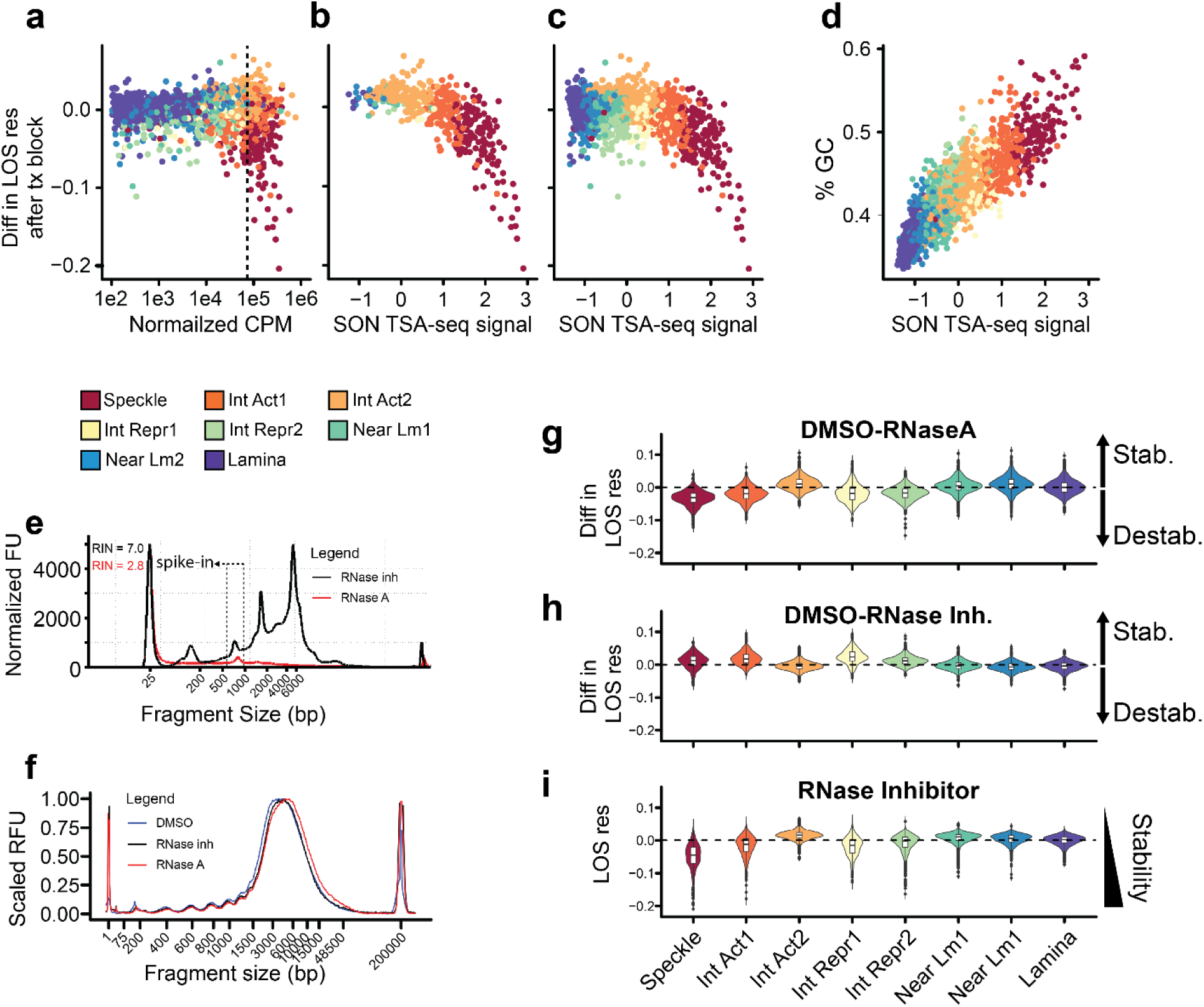
RNA abundance and proximity to speckle dictate chromatin interaction stability at speckles. **a**, Scatter plot showing difference in LOS residuals after transcription block against nascent RNA mapped to these regions binned at 1 Mb and colored by SPIN state. Scatter plots showing difference in LOS residuals after transcription block against SON-TSA-seq signal associated with these regions for regions with the highest RNA output [**b**,regions to the right of the dotted line in (a)] and all genomic regions (**c**). **d**, scatter plot showing the relationship between GC content and SON-TSA-seq signal. **e**, Fragment analyzer profile showing degradation of RNA after RNaseA treatment. **f**, Fragment analyzer profiles of DNA fragments resulting from LC-Hi-C in situ pre-digestion of unfixed nuclei isolated treated with RNaseA, DMSO or preserved with RNase inhibitor. Global difference between control and RNase-treated LOS residuals(**g**) and between control and RNase inhibitor LOS residuals (**h**) for all SPIN states. **i**, LOS residuals for LC-Hi-C experiments where RNase inhibitors were used during nuclei isolation and DpnII pre-digestion. Bootstrap analysis generated significant p values (p < 0.0001) for comparisons of each SPIN group to 0. All boxplots show the median (center line), interquartile range (box), and 1.5×IQR whiskers, with outliers plotted individually. Scatter plots are binned at 1 Mb and violin plots are binned at 150 kb.

**Extended Data Fig. 7:**
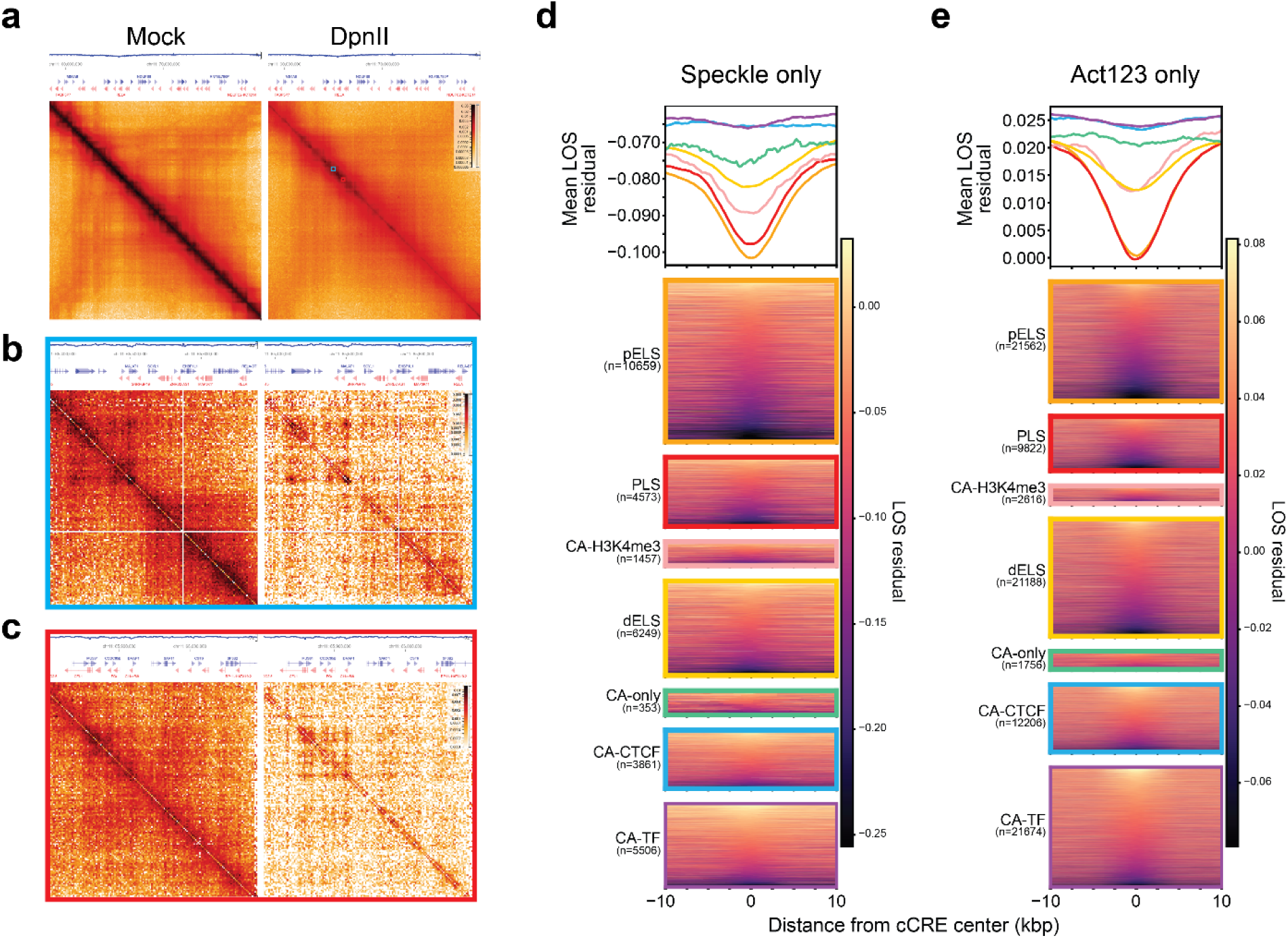
**a**, Interaction frequency map binned at 25 kb for Mock-and DpnII-digested LC-Hi-C libraries with associated LOS track and gene annotations for positions 60-75 Mb along chromosome 11. Zoom in of position 65.4-65.7 Mb (**b**) and position 65.8-66.1 Mb regions (**c**) both binned at 2kb. LOS residuals (binned at 5kb) centered on cCREs in either speckle-(**d**) or Int_Act123 (**e**) regions are shown individually and averaged by 200bp bins across the region.

