## Supplementary Figures for "Formation of an RNA-mediated nuclear compartment"

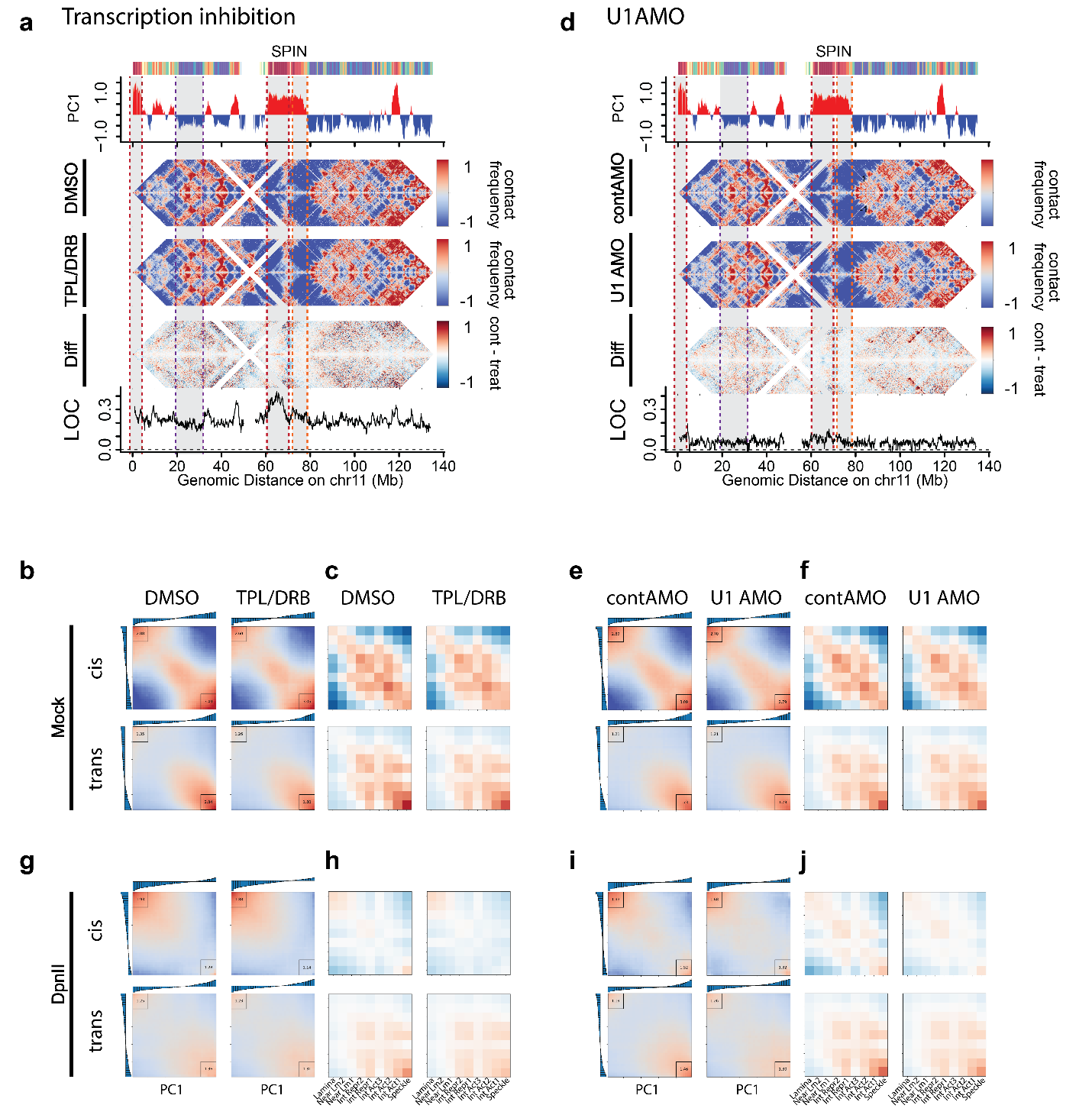
 **Supplementary Fig. 1: a**, Tracks for PC1, SPIN annotation and LOC (see Methods) along with log2(obs/exp) heatmaps of chromosome 11 for control and treated Mock-digested libraries for TPL/DRB (**a**) and  U1AMO (**d**) treatments. Cis and trans PC1 saddle plot for TPL/DRB (**b**) and  U1AMO (**e**) treatments. Cis and trans saddle plots categorized by SPIN state for TPL/DRB (**c**) and  U1AMO (**f**) treatments. Associated cis and trans PC1 saddles (**g** and **i**), SPIN-categorized saddles (**h** and **j**) generated from the DpnII-digested LC-Hi-C libraries for TPL/DRB and  U1AMO treatments. Heatmaps are binned at 250 Mb and tracks are binned at 50 kb.

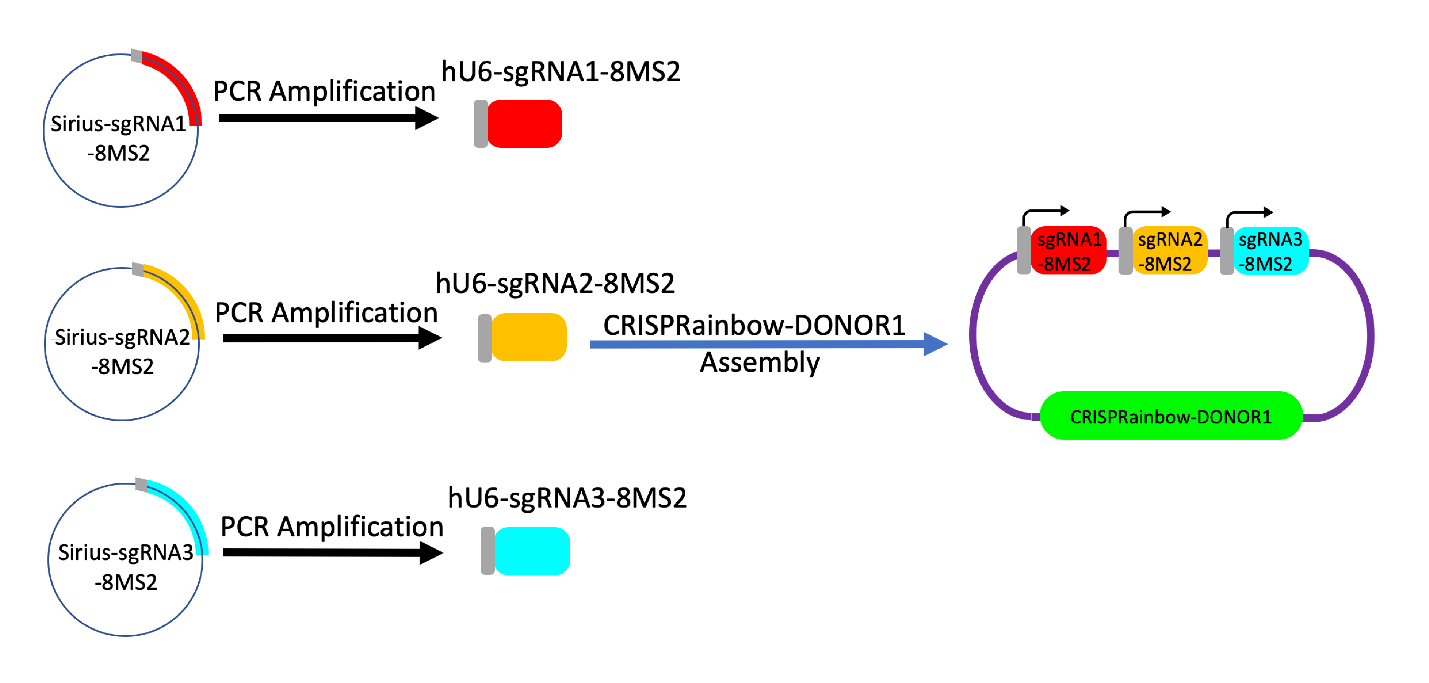
 **Supplementary Fig. 2:** Assembly of one plasmid for the expression of multiple CRISPR-Sirius-sgRNAs. The schematic illustrates the steps for constructing an assembled 3x single-guide RNA (sgRNA) plasmid. Individual sgRNA cassettes with human U6 (hU6) promoter from the CRISPR-Sirius-sgRNA expression vector, pPUR-hU6-sgRNA-Sirius-8XMS2 (Addgene #121942), are amplified by PCR and assembled into the CRISPRainbow-DONOR1 vector (Addgene #75398) using a one-step Gibson assembly reaction. For the C19-Stable locus, sgRNA1 is C19-d01, sgRNA2 is C19-d05, and sgRNA3 is C19-d06. For the C19-Destabilized locus, sgRNA1 is C19-G03 and sgRNA2 is C19-22; only two gRNAs are assembled for this construct.

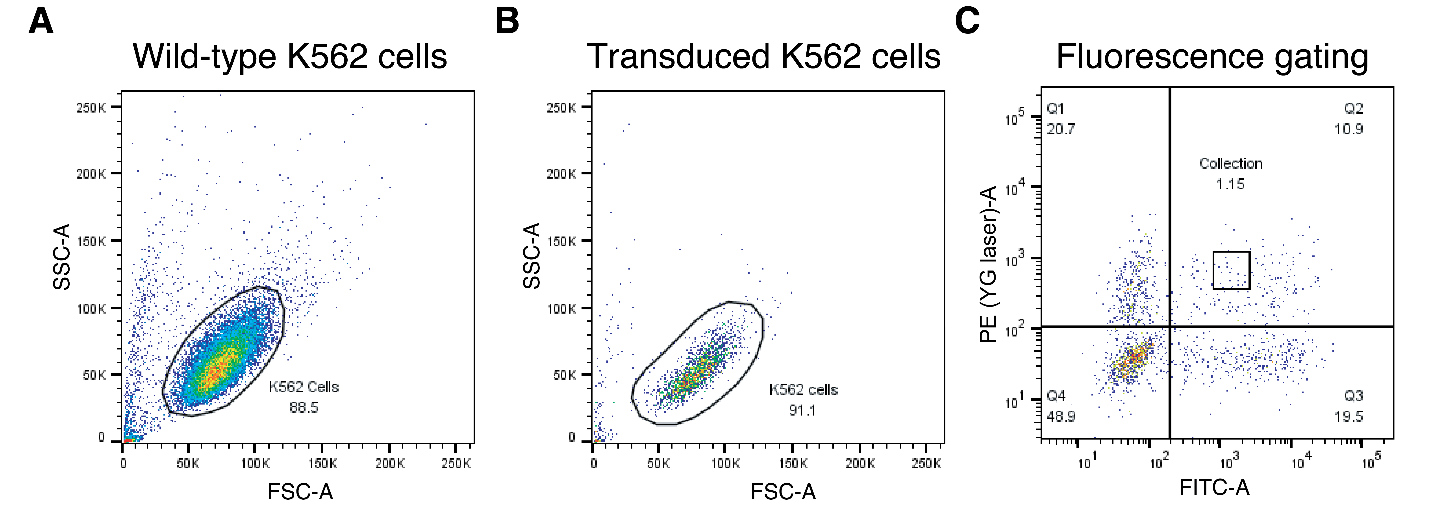
 **Supplementary Fig. 3:** Enrichment of fluorescent cells by flow cytometry sorting. **a**, K562 wild-type (non-fluorescent) cells used for size and granularity gating (FSC/SSC). **b**, K562 cells following lentiviral transduction with MCP-tdTomato and PCP-GFP. **c**, Representative flow cytometry plots showing fluorescence gating in the PE (YG laser, 561 nm) and FITC (blue laser, 488 nm) channels. Quadrant percentages are as follows: double-negative (48.9%), tdTomato-positive (20.7%), GFP-positive (19.5%), and double-positive (10.9%). Gating was established to exclude cells with extremely high or low fluorescence intensity (1.15%). Although PCP-GFP was initially gated in the FITC-A channel, this marker was excluded from the final study due to a significant decrease in GFP signal during post-sort culture. Figures were generated from 2,000–20,000 events per sample.

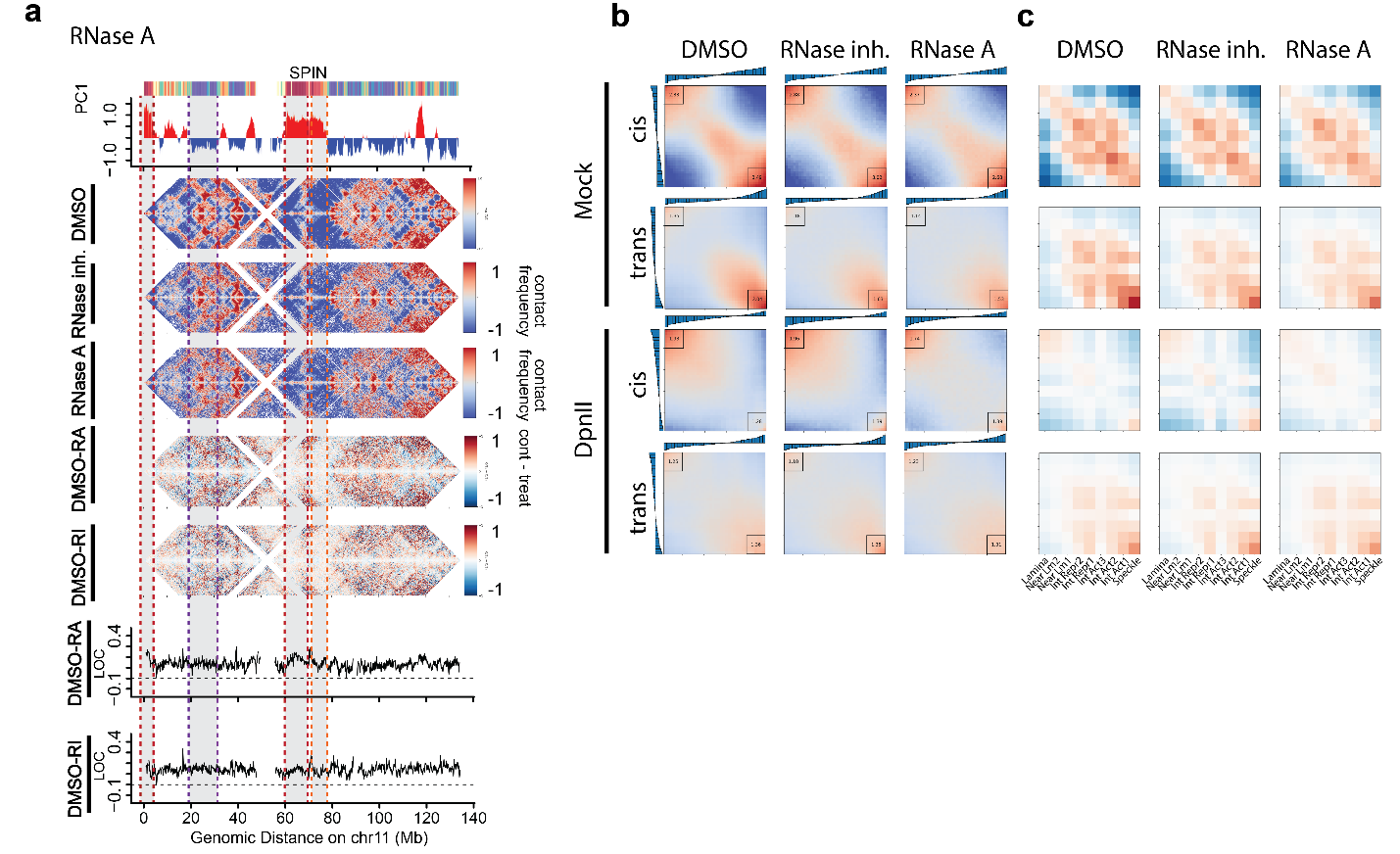
 **Supplementary Fig. 4:** Tracks for PC1, SPIN annotation and LOC (see Methods) along with log2(obs/exp) heatmaps of chromosome 11 for control and treated libraries for RNaseA treatment and associated control (**a**). Cis and trans PC1 saddle plots (**b**) along with saddle plots categorized by SPIN state (**c**) generated from either the RNaseA treatment or controls for Mock- or DpnII-digested libraries.  Heatmaps are binned at 250 Mb and tracks are binned at 50 kb.
